# Language-guided domain adaptation improves protein sequence representations for antimicrobial resistance analysis

**DOI:** 10.64898/2026.11.04.764426

**Authors:** Rongye Ye, Lun Li, Zongyi Geng, Shuhui Song

**Author notes:** To whom correspondence should be addressed: **Contact:** (Song S).

## Abstract

Antimicrobial resistance (AMR) in bacterial pathogens complicates infectious disease management. Protein sequences encode molecular features underlying resistance functions such as antibiotic inactivation and drug efflux, providing a basis for investigating bacterial AMR. Protein language models can extract transferable representations from these sequences. However, general-purpose protein language models have limited capacity to represent AMR-associated sequences, while task-specific tools often address individual prediction problems. These limitations highlight the need for protein representations that better capture resistance-associated functions and can be reused across complementary tasks, supporting systematic AMR analysis from sequence variation to functional relationships among proteins. We developed BacLLM, a language-guided protein representation framework that integrates annotation-derived functional knowledge into protein language model representations for AMR analysis in bacterial pathogens. Through sequence enhancement, semantic enrichment of UniProt annotations, and cross-modal contrastive learning, BacLLM transfers LLM-encoded annotation priors into the sequence encoder, enabling sequence-only inference after training. Compared with baseline models, BacLLM provided more informative representations of resistance-associated sequences, achieving a mean F1 score of 0.956 across six AMR benchmarks with logistic regression, 2.7 percentage points above the strongest baseline. It also captured resistance-associated mutational signals at single-residue resolution and preserved taxonomic relationships not reflected by phenotype-based clinical groupings. Moreover, BacLLM recovered resistance-associated functional organization within a TolC-centered protein association network, prioritizing proteins with low sequence identity to TolC but predicted structural similarity to TolC. Extending the same representation framework to an intervention-oriented application, BacLLM also improved strain-level phage–host matching for phage therapy candidate prioritization. Together, these findings establish BacLLM as a reusable protein representation framework for analyzing resistance-associated sequence variation and functional organization, with applications in AMR surveillance and phage candidate prioritization.

## Introduction

Antimicrobial resistance (AMR) in bacterial pathogens continues to emerge and spread, posing a major global public health threat [1,2]. In 2019, bacterial AMR was estimated to directly cause approximately 1.27 million deaths and to be associated with nearly 5 million deaths worldwide [3]. In response to this growing burden, the World Health Organization released an updated bacterial priority pathogens list in 2024, emphasizing the need for improved infection control, resistance surveillance, and new antimicrobial interventions [4]. Bacterial resistance is often mediated by protein functions, including antibiotic inactivation and efflux. Protein sequence analysis provides molecular-level information for AMR surveillance and resistance mechanism characterization [5,6]. In addition, protein-level determinants contribute to phage–host recognition, linking sequence analysis to phage therapy candidate prioritization.

Although experimental assays remain the gold standard for AMR-related analyses, their cost, time requirements, and limited throughput constrain the analysis of rapidly expanding microbial sequence data. To address these limitations, computational methods have traditionally inferred resistance-associated protein functions through sequence similarity-based methods. However, when query proteins have low similarity to annotated references, alignment-based methods can produce substantial false negatives [7,8]. More recently, task-specific deep learning models have improved alignment-free prediction in selected AMR-related applications. For example, ARGNet combines autoencoders and convolutional neural networks for resistance gene detection [7], HMD-ARG uses hierarchical multitask learning to predict resistance categories, mechanisms, and gene mobility [8], and PhageHostLearn models receptor-binding proteins and host receptors for strain-level phage–host prediction in *Klebsiella pneumoniae* [9]. Although these models have achieved promising performance in their respective tasks, they often depend on specialized architecture or task-specific features [10], limiting their reuse across the interconnected analyses required for AMR surveillance, mechanistic investigation, and therapeutic prioritization.

Pretrained protein language models offer an alternative by learning transferable representations from large-scale unlabeled sequences [11–13]. General-purpose models trained on panspecies data include sequence-only protein language models such as ESM-2 [14] and multimodal models such as ProTrek [15]. Although these models capture broad evolutionary, biophysical, structural, and functional patterns, the broad taxonomic coverage of their training data may underrepresent AMR-relevant signals. Sequence-only models such as ESM-2 face an additional modality limitation because their representations are learned exclusively from amino acid sequence context, without explicit semantic supervision from functional annotations or AMR-relevant biomedical knowledge. ProTrek alleviates this modality limitation by aligning protein sequences with structural and textual information, but its general-purpose training does not specifically resolve the shared AMR domain mismatch. Bacteria-focused models such as Bacformer [16] partially address the domain gap by incorporating bacterial context through complete-proteome representations; however, treating the complete proteome as an integrated unit may dilute fine-grained signals from individual proteins and reduce sensitivity to functionally important sequence variation. Collectively, these models do not simultaneously combine AMR-specific adaptation, explicit integration of biomedical knowledge, and protein-level granularity. These capabilities are important because digital AMR analysis spans multiple levels of protein sequence interpretation. Sequence-based identification and functional characterization of antimicrobial resistance genes (ARGs) support AMR surveillance and resistance mechanism analysis [17]. Mutation-effect estimation characterizes the resistance-associated consequences of local sequence changes [18], whereas protein functional network analysis captures functional relationships among proteins. For therapeutic prioritization, matching phage receptor-binding protein sets with bacterial receptor-associated protein sets enables the selection of candidate phages for specific bacterial strains [9]. Together, these applications require a domain-adapted, annotation-informed protein representation that preserves fine-grained sequence information while capturing functional relationships among multiple proteins.

To address these representation requirements, we developed BacLLM, a language-guided protein representation model for AMR analysis in WHO priority bacterial pathogens. BacLLM was trained on approximately 12,000 curated sequence-annotation pairs from UniProt, covering these pathogens. To facilitate the transfer of annotation-derived knowledge into protein sequence representations, we designed complementary sequence enhancement and annotation semantic enhancement modules. On the sequence side, a learnable sequence enhancement module refines protein embeddings produced by ESM-2 and projects them into a shared sequence-text semantic space. On the text side, the annotation semantic enrichment module organizes the information contained in UniProt annotations into context-rich biomedical descriptions, which are subsequently encoded by Qwen3 [19] into dense semantic representations. Through domain-adaptive cross-modal contrastive learning, BacLLM aligns enhanced protein sequence embeddings with annotation-derived semantic representations, thereby incorporating biomedical annotation priors into the protein sequence representation space. After training, the text branch is removed, enabling sequence-only inference for downstream applications.

We evaluated BacLLM across multiple complementary applications involving AMR sequence classification, single-residue mutation analysis, protein functional network analysis, and strain-resolved phage–host matching based on receptor-associated protein sets. Across benchmark AMR prediction tasks, BacLLM provided more informative protein sequence representations than mainstream protein language models. Beyond classification, BacLLM assessed resistance-associated effects of single-residue mutations through in silico mutagenesis, using changes in ARG prediction probabilities. Within a TolC-centered functional association network constructed from its representations, BacLLM further prioritized proteins with low sequence similarity to TolC but predicted structural similarity to TolC. BacLLM also improved the prioritization of candidate phages for specific bacterial strains through phage–host interaction prediction. Together, these findings support BacLLM as a protein sequence representation framework for digital AMR surveillance, resistance mechanism investigation, and phage therapy candidate prioritization.

## Materials and Methods

### Overall structure

To adapt general protein language model representations to WHO priority bacterial pathogens associated with AMR, we developed BacLLM, a domain-adaptive multimodal contrastive learning framework (Fig. 1). BacLLM adopts an asymmetric dual-encoder architecture that aligns bacterial protein sequences with their paired UniProt annotations, thereby transferring annotation-derived biomedical priors into the protein sequence representation space.

**Fig. 1.**
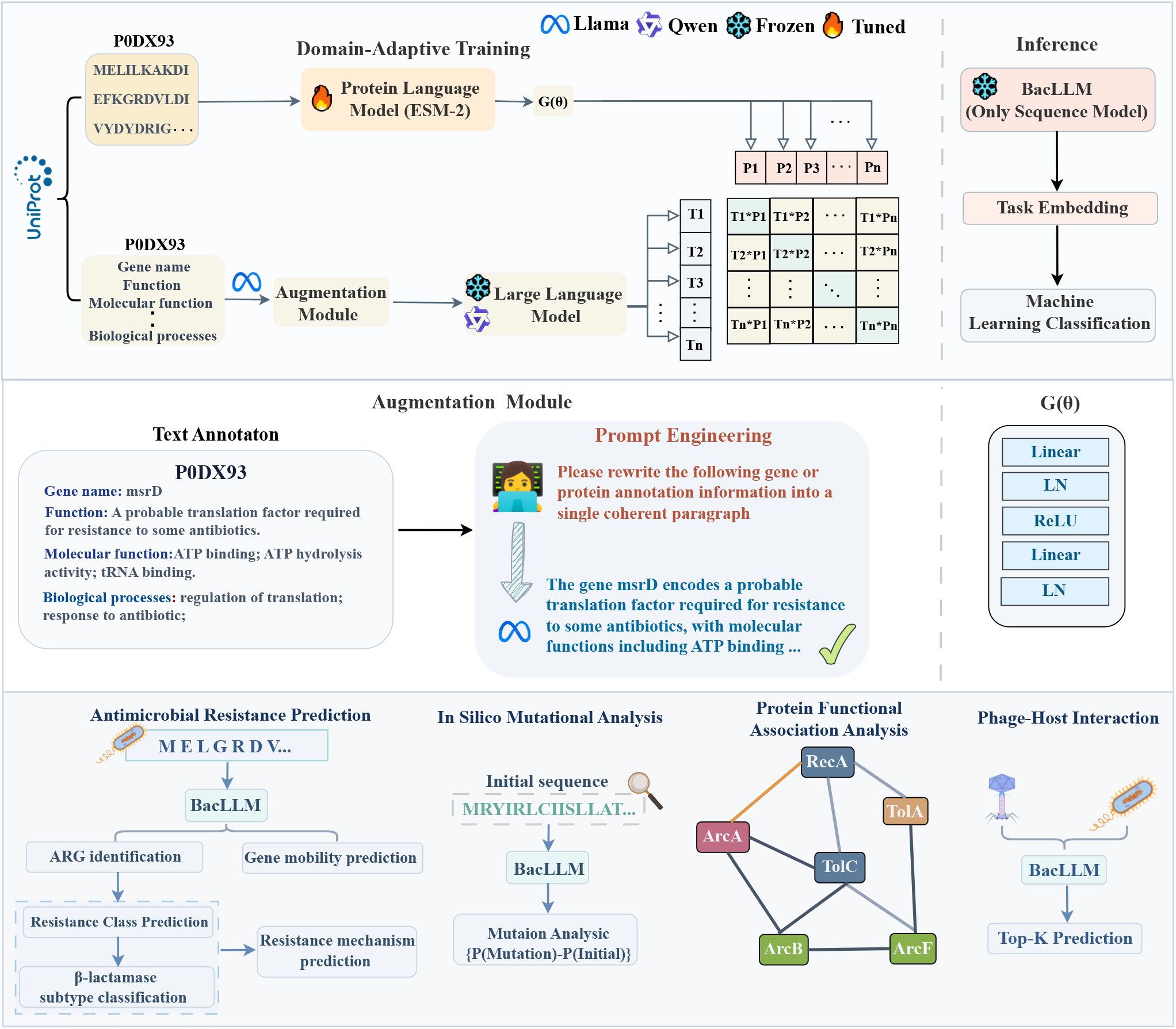
Overview of the BacLLM architecture. The proposed framework consists of three core components: a multimodal contrastive-learning pipeline for training and inference; sequence and text semantic enhancement modules; and downstream analytical and clinical applications enabled by BacLLM representations.

The framework consists of two coordinated branches. In the sequence branch, protein sequences from bac-terial pathogens included in the WHO priority pathogens list are encoded by ESM-2, which serves as the trainable foundation sequence encoder. To improve domain adaptation and cross-modal alignment, we designed a learnable sequence enhancement module, *G(θ),* that refines protein embeddings produced by ESM-2 and projects them into a shared sequence-text semantic space. This module is designed to enhance AMR-associated functional signals in protein sequence representations and project the resulting representations into a shared latent space aligned with the target text representations.

In the text branch, the input consists of UniProt text annotations paired with protein sequences. To enrich the semantic content of these annotations, we designed a semantic enhancement module that transforms discrete biological knowledge into coherent and context-rich natural language descriptions through prompt engineering. The reconstructed text is then processed by the Qwen3 model to extract dense textual representations containing expert prior knowledge. During joint training, matched sequence-text pairs are optimized using a contrastive loss function that pulls each protein embedding toward the semantic representation of its paired annotation while separating it from mismatched annotations within the same batch. Through this process, BacLLM internalizes UniProt-derived biomedical priors into the trainable sequence encoder. After training, the text branch is removed, and BacLLM performs sequence-only inference: users provide raw amino acid sequences, and the model outputs function-aware protein representations for down-stream AMR-related analyses.

To enable foundation sequence models to absorb complex human prior knowledge, we constructed a multimodal contrastive learning framework based on an asymmetric dual-encoder architecture. Given a training dataset 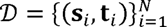 containing *N* samples, where *s_i_* denotes an input protein sequence and *t_i_* denotes its paired textual prior annotation, the objective is to learn a cross-modal joint representation space. During training, a batch of size s is sampled for optimization. Given a natural language description *t_i_*, we employ Qwen3 as the text encoder *h_ψ_(·).* Since large language models have absorbed extensive knowledge and possess highly structured semantic spaces, the parameters *ψ* are frozen throughout training. The input text is first tokenized and then processed by the text encoder to obtain the final-layer hidden states. Mean pooling is subsequently applied across all valid tokens, followed by *L_2_* normalization:

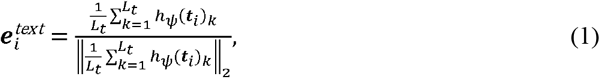

where *L_t_* denotes the number of valid tokens in the input text sequence. In the sequence encoding branch, we employ ESM-2 as the foundation sequence model, denoted as *f_e_(·),* with all parameters *fJ* remaining trainable throughout optimization. Given an input protein sequence *s_i_*, the sequence is encoded and subsequently mean pooled to produce an initial high-dimensional representation, 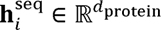. To bridge both the dimensional discrepancy and semantic gap between the sequence and text modalities, the sequence representation is further processed by a semantic enhancement module, *G(θ).* This module consists of two fully connected layers combined with layer normalization to maintain numerical stability during training. The transformation is defined as:

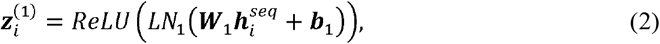

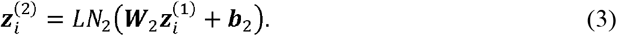

Here, *LN_1_* and *LN_2_* denote layer normalization. The projected sequence representation is subsequently normalized using *L_2_* normalization to obtain the final protein embedding in the aligned space:

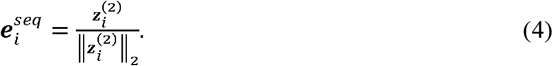

Through this mapping process, the latent protein representation is explicitly projected into a *d_text_* dimensional joint latent space aligned with the text representations. Since our objective is to inject knowledge from a static text semantic space into a dynamically updated sequence model, we adopt a unidirectional InfoNCE loss. Given a batch of size s, each paired sequence and text embedding 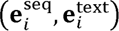 is treated as a positive pair, while all other mismatched combinations within the batch are regarded as negative pairs. The objective is computed only in the sequence-to-text direction:

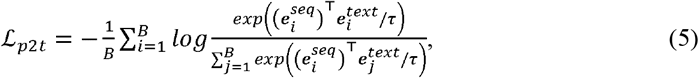

where *T* is a temperature parameter controlling the smoothness of the probability distribution. Since the text encoder is frozen during training, we adopt a unidirectional objective. This objective encourages the trainable ESM-2 encoder to progressively align its representation space with the semantic space encoded by Qwen3.

## Dataset

### Domain-Adaptive Training Dataset

To support domain-adaptive cross-modal training, we constructed a paired sequence-annotation dataset from clinically important bacterial pathogens prioritized by the WHO. The dataset focused on WHO priority bacterial pathogens (Supplementary Table S1) associated with AMR and contained approximately 12,000 protein sequence-annotation pairs curated from UniProt [20]. For each protein entry, we extracted available annotation fields, including gene names, protein products, Gene Ontology (GO) terms, and functional descriptions. Because UniProt annotations are often stored as discrete structured fields, they may lack the contextual continuity required for effective text-sequence alignment. We therefore designed an annotation semantic enhancement strategy to reorganize these curated fields into coherent biomedical text while preserving the original UniProt-derived information. Specifically, MedLLM-Llama3-8B was used to integrate the extracted annotation fields into context-rich descriptions through predefined prompts (Supplementary Fig. S1). The enhanced annotation text was then paired with the corresponding protein sequence to form the final training dataset for domain-adaptive multimodal contrastive learning.

### Antimicrobial Resistance Prediction Datasets

To evaluate BacLLM across the major stages of AMR sequence analysis, we organized the benchmark datasets into three complementary objectives: ARG detection, functional characterization of resistance, and gene mobility prediction. Unless otherwise specified, all benchmark datasets were randomly split into training and test sets at a ratio of 8:2. To assess the robustness of the model comparison, we additionally performed a targeted MMseqs2-guided, sequence-identity-controlled five-fold cross-validation analysis for the first downstream application, which comprised six AMR sequence-prediction benchmark settings spanning binary and multiclass endpoints. MMseqs2 was used to cluster protein sequences at minimum sequence-identity thresholds of 85%, 90%, 95%, and 98%, with 80% bidirectional sequence coverage (-c 0.8, --cov-mode 0) and connected-component clustering (--cluster-mode 1). At each threshold, all sequences belonging to the same MMseqs2 cluster were assigned to the same fold, preventing members of that cluster from occurring in both the training and test partitions. This analysis was used to determine whether the comparative performance of the protein representations remained robust under different sequence-identity constraints.

### ARG detection dataset

The first benchmark aimed to evaluate whether an unknown amino acid sequence could be identified as an antimicrobial resistance gene (ARG). Positive samples comprised approximately 27,000 annotated resistance protein sequences collected from the ARG reference database ARGNet-DB [7]. To construct the negative set, we collected protein sequences from bacterial pathogens included in the WHO priority pathogens list using the NCBI database. Given the lack of a comprehensive and well-annotated non-ARG reference database, we drew on the negative sampling strategy employed in HMD-ARG and screened these sequences against ARGNet-DB using BLAST. To construct a challenging negative set and reduce shortcut learning from trivially dissimilar sequences, we retained candidate proteins with alignment coverage exceeding 50% and sequence identity ranging from 50% to 95% relative to reference ARGs, and an E-value ≤ 1 x 10^⁻⁵^. Sequences with identity greater than 95% were excluded to reduce the likelihood of including proteins highly similar to known ARGs.

### Functional characterization of resistance dataset

After ARG detection, we evaluated whether resistance-associated sequences could be assigned more specific functional annotations. Resistance class and mechanism prediction datasets were constructed from ARGNet-DB [7] and HMD-ARG [8]. For resistance class prediction, HMD-ARG and ARGNet-DB originally contained 33 and 36 classes, respectively. Classes containing fewer than 20 sequences were consolidated into an “other” class to reduce extreme class imbalance, resulting in 18 classes for HMD-ARG and 22 classes for ARGNet-DB (Supplementary Fig. S2a, b). For resistance mechanism prediction, we used the seven major resistance mechanisms annotated in HMD-ARG and applied the same consolidation strategy to mechanism categories containing fewer than 20 sequences (Supplementary Fig. S2c). We further evaluated fine-grained discrimination of clinically important resistance enzymes using the Beta-Lactamase Database (BLDB) [21]. After data curation, six major β-lactamase subtypes were retained for downstream classification (Supplementary Fig. S2d).

### Gene mobility prediction dataset

To evaluate the ability of BacLLM to distinguish between intrinsic and acquired resistance genes, we constructed a gene mobility prediction benchmark using gene mobility annotations from HMD-ARG.

### Protein Functional Association Datasets

To evaluate whether BacLLM captures functional relationships between proteins in clinically important bacteria, we constructed protein functional association benchmark datasets using data from STRING [22], a comprehensive database of known and predicted protein–protein associations. Given the scale of STRING, we restricted the analysis to the six clinically important ESKAPE bacteria (Supplementary Table S2) [23], which are frequently associated with difficult-to-treat infections and antimicrobial resistance in clinical settings. We focused on three STRING evidence channels or scores: Neighborhood (conserved genomic proximity), Co-occurrence (phylogenetic co-occurrence across genomes), and Combined Score (integrated confidence from multiple evidence sources). To convert continuous STRING confidence scores into binary labels for model training, we followed the threshold used in previous studies [16]: protein pairs with scores greater than or equal to 600 were labeled as interacting, whereas pairs with scores below 600 were labeled as non-interacting. To reduce evaluation bias from duplicated protein-pair records, all datasets underwent strict symmetric deduplication. Detailed dataset statistics are provided in Supplementary Table S2. For model input construction, sequence embeddings were extracted independently for the two proteins in each pair and concatenated along the feature dimension. The concatenated embeddings were then used together with the corresponding binary labels for downstream prediction. Each dataset was randomly divided into training and test sets at a ratio of 8:2.

### Phage–Host Interaction Datasets

To evaluate BacLLM in a precision antibacterial intervention setting, we constructed phage–host interaction prediction benchmarks using two experimentally validated strain-level datasets. Because clinical phage therapy depends on matching candidate phages to bacterial strains [24], all interaction labels used in this study were defined at the strain level.

First, we used a Klebsiella pneumoniae phage–host interaction dataset [9] comprising 105 genetically diverse phage genomes and 200 bacterial genomes. This dataset contains 10,006 spottest results, including 333 experimentally confirmed positive phage–host interactions. A visualization of a subset of the dataset is provided in Supplementary Fig. S3a. For phage feature extraction, we followed the original study [9] and used receptor-binding proteins (RBPs) as the primary phage-side features. On the bacterial side, the capsular polysaccharide synthesis locus (K-locus) was extracted as a host receptor-associated feature. Experimentally validated spottest results were used as ground truth labels. For evaluation, bacterial strains were clustered using the Bidirectional Best Match algorithm (Supplementary Note S1), and leave-one-cluster-out cross-validation (LOCO-CV) was used for model training and performance assessment.

Second, we introduced an Escherichia dataset [25] as an additional benchmark to evaluate the cross-strain representation capability of the model. This dataset consists of 403 natural isolates representing the evolutionary diversity of the genus and 96 phages targeting these isolates, resulting in 38,688 recorded interactions. The original dataset quantified interaction strength using the Minimum Lytic Concentration (MLC) score, defined as the lowest phage concentration at which a lytic interaction was observed. Since the primary objective was to predict whether an interaction occurs between each phage–host pair, the interaction labels were binarized. All infection events were labeled as positive interactions, whereas non-infection events were labeled as non-interactions. A visualization of a subset of the dataset is provided in Supplementary Fig. S3b. For feature extraction, RBPs were again used as the primary phage features. On the bacterial side, phage recognition in Escherichia species is characterized by a highly complex and cooperative interaction pattern rather than being governed by a single dominant surface antigen. Previous studies have demonstrated that outer membrane proteins (OMPs) [26,27], lipopolysaccharides (LPS), and capsular poly-saccharides [28,29] all play critical roles in mediating phage infection. Therefore, OMPs together with key proteins involved in LPS and capsule biosynthesis and secretion were extracted to construct a composite set of host receptor-associated features. Given the large scale of this dataset, the computational cost of LOCO-CV was prohibitively high. Therefore, the dataset was randomly divided into training and test sets at a ratio of 8:2. This procedure was repeated 20 times using different random seeds, and the final results were reported as the average across all runs.

### Baseline Setting

To evaluate the representation capability of BacLLM, we compared it with four representative protein language models (PLMs) widely used in protein sequence analysis. All pretrained models were evaluated under a standardized frozen-encoder setting. Specifically, each model was used as a fixed feature extractor to obtain protein embeddings, which were then used as inputs to relatively simple downstream classifiers. This frozen-encoder evaluation strategy was designed to assess the discriminative quality of the embeddings learned by each pretrained model while limiting task-specific model complexity and avoiding fine-tuning of the protein encoders [30–32].

The baseline models included Ankh [33], a lightweight general-purpose protein language model designed to achieve strong representation performance with reduced model size and computational cost; ESM-2 [14], a widely used protein language model pretrained on large-scale pan-species protein corpora, for which we selected the 33M-parameter version to match practical deployment constraints; ProteinBERT [34], a protein language model designed for protein property prediction and efficient processing of long protein sequences; and ProTrek [15], a multimodal protein language model that aligns amino acid sequences, protein structures, and natural language descriptions in a shared embedding space. In addition to these baselines, we evaluated two BacLLM variants developed in this study. BacLLM (4B) uses ESM-2 (33M) as the sequence encoder and Qwen3-4B as the text encoder during multimodal training, whereas BacLLM (8B) uses the same ESM-2 (33M) sequence encoder with Qwen3-8B as the text encoder. Because the text branch is removed after training, both BacLLM variants have the same sequence encoder size during deployment.

For downstream prediction, we employed logistic regression (LR) as a linear classifier and support vector machines with a radial basis function (RBF) kernel as a relatively simple nonlinear classifier. These classifiers enabled the evaluation of both linear separability and nonlinear discriminability of the extracted protein embeddings, providing a consistent comparison of representation quality across different PLMs without introducing complex task-specific neural architectures.

## Results

### BacLLM enhances antimicrobial resistance sequence representation

To evaluate whether BacLLM supports a progressive sequence-based analysis of AMR, we assessed five biological prediction tasks across six primary benchmark settings using a standardized frozen-encoder evaluation protocol with lightweight linear and relatively simple nonlinear classifiers. BacLLM was compared with four representative protein language models, including Ankh, ESM-2, ProteinBERT, and ProTrek, using frozen sequence representations and identical LR and SVM classifiers. The five biological tasks comprised ARG identification, gene mobility prediction, antibiotic resistance class prediction (evaluated independently on ARGNet-DB and HMD-ARG), β-lactamase subtype classification, and resistance mechanism prediction. We additionally evaluated model performance on low-frequency resistance mechanism labels.

We first evaluated whether BacLLM representations could distinguish ARGs from non-ARG sequences, which is the initial step in sequence-based AMR analysis (Fig. 2a). With SVM, BacLLM (4B) and BacLLM (8B) achieved F1 scores of 0.936 and 0.940, respectively, outperforming all baseline models. With LR, ProteinBERT achieved a slightly higher F1 score than BacLLM (8B) (0.920 versus 0.916). BacLLM remained competitive with the best-performing baseline. These results establish that BacLLM preserves the sequence-level information required for ARG discrimination.

**Fig. 2.**
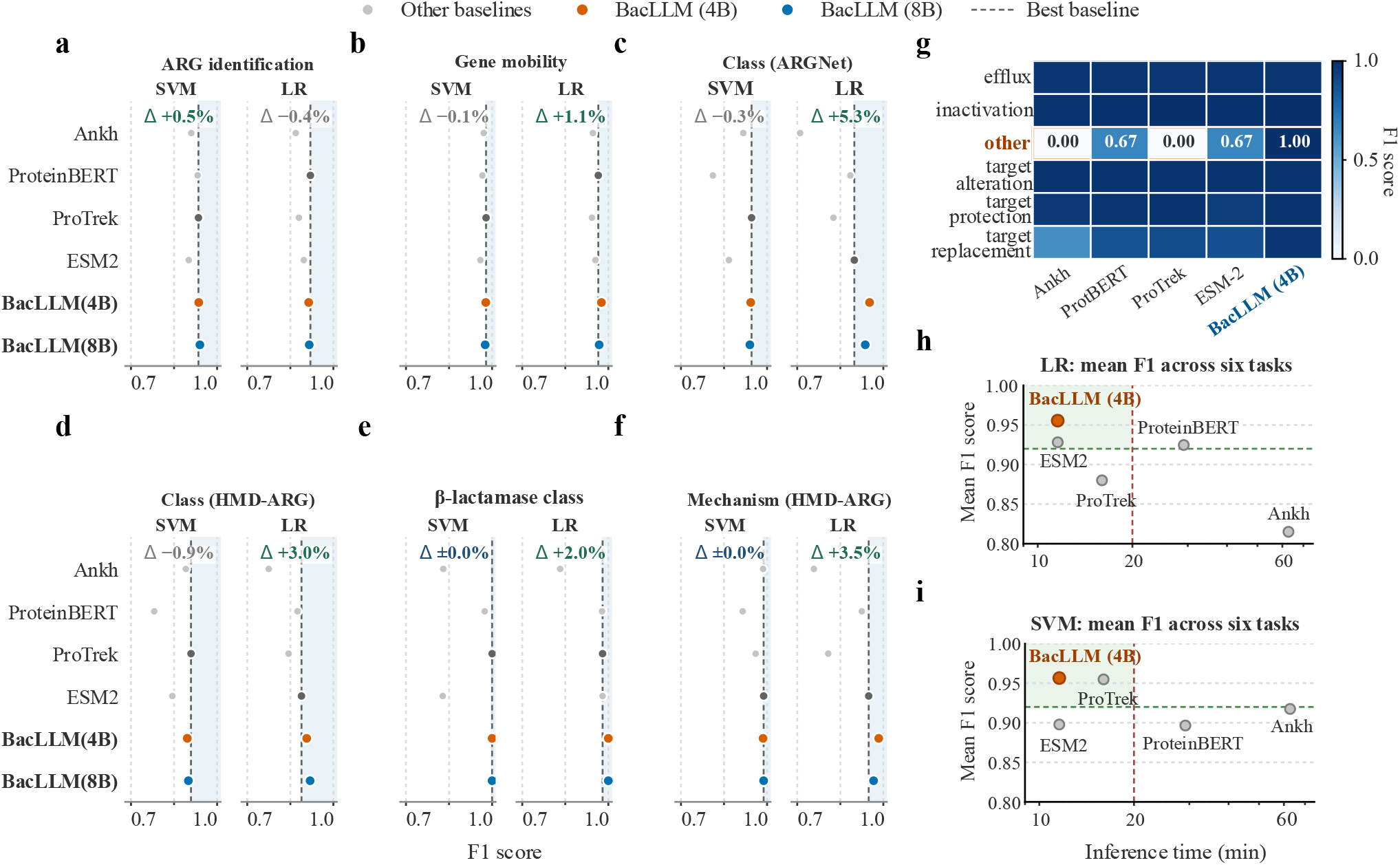
Benchmark performance of BacLLM across antimicrobial resistance prediction tasks. **a**, ARG identification task, i.e., binary classification to predict whether unknown sequences are ARGs; **b**, gene mobility prediction for distinguishing between endogenous and exogenous (horizontally transferable) ARGs; **c**, resistance category prediction performance based on the ARGNet database; **d**, resistance category prediction performance evaluated on the HMD-ARG dataset; **e**, β-lactamase class prediction performance evaluated on the BLDB database; **f**, resistance mechanism prediction performance evaluated on the HMD-ARG dataset; **g**, few-shot learning capability analysis of BacLLM on the HMD-ARG dataset; **h, i**, trade-off analysis between embedding extraction time and mean F1 score across six benchmark tasks for each model, using LR (h) and SVM (i) as downstream classifiers.

We next examined whether BacLLM could predict ARG mobility, as horizontal gene transfer facilitates the spread of resistance determinants across bacterial populations, making gene mobility an important consideration in AMR surveillance (Fig. 2b). BacLLM (4B) with LR achieved the highest F1 score of 0.976, outperforming ProTrek by 3.3 percentage points. With SVM, BacLLM (4B) and ProTrek performed similarly, differing by only 0.1 percentage points. Thus, BacLLM representations captured information relevant not only to ARG identity but also to their potential mobility.

To determine which antibiotic classes were associated with the identified resistance sequences, we subsequently evaluated resistance class prediction on ARGNet-DB and HMD-ARG. With LR, BacLLM (4B) improved the F1 score over ESM-2 by 5.3 percentage points on the ARGNet-DB dataset (Fig. 2c). On the HMD-ARG dataset, BacLLM (4B) and BacLLM (8B) outperformed ESM-2 by 1.8 and 3.0 percentage points, respectively, and ProTrek by 6.3 and 7.5 percentage points, respectively (Fig. 2d). Although ProTrek outperformed BacLLM (8B) by 0.9 percentage points with SVM, the stronger LR performance of both BacLLM variants indicates that their representations support clearer linear discrimination of resistance classes. The consistent results across two databases support the robustness of BacLLM for antibiotic resistance class characterization.

Building on broad resistance class prediction, we next examined whether BacLLM could resolve finer functional distinctions within β-lactamases, a clinically important group of resistance enzymes. In the β-lactamase subtype classification benchmark, both BacLLM variants achieved an F1 score of 1.0 (Fig. 2e), demonstrating strong discrimination among the evaluated subtypes. This result extends the evaluation from broad antibiotic resistance classes to more specific functional variation within a major resistance-enzyme family.

We next investigated whether BacLLM could distinguish the mechanisms underlying antibiotic resistance, rather than merely identifying the associated antibiotic classes. On the HMD-ARG resistance mechanism dataset, BacLLM (4B) achieved the highest LR F1 score of 0.984, outperforming ESM-2, the best-performing baseline, by 3.5 percentage points (Fig. 2f). To further assess its representation capacity when labeled examples were limited, we examined the low-frequency “other” label, which contained only 20 samples. BacLLM retained a clear performance advantage with LR in this setting (Fig. 2g). Together, these results indicate that BacLLM representations encode information relevant to distinct resistance mechanisms and remain discriminative for low-frequency labels, extending their utility from ARG identification and classification to resistance mechanism characterization.

We then summarized overall performance by averaging F1 scores across the six primary benchmark settings and comparing them with measured protein embedding extraction time (Fig. 2h, i). At the individual-setting level, at least one of the two BacLLM variants achieved the highest F1 score in five of the six settings with LR. With SVM, at least one BacLLM variant ranked first or tied for first in three settings and ranked second in the remaining three, with only marginal differences from the best-performing baseline (Supplementary Tables S3–S4). Precision and recall showed trends consistent with the F1 results (Supplementary Table S3). BacLLM (4B) achieved higher mean F1 scores than all baseline models under both classifiers while maintaining rapid embedding extraction.

Finally, to examine the robustness of the model comparison under different sequence-identity constraints, we performed additional MMseqs2-guided five-fold cross-validation at sequence-identity thresholds of 85%, 90%, 95%, and 98% across the six AMR prediction benchmark settings (Supplementary Fig. S4). For each model and threshold, F1 scores were first averaged across five folds within each benchmark setting and then averaged equally across the six settings. With LR, BacLLM (8B) ranked first at all four thresholds, achieving mean F1 scores of 0.877, 0.889, 0.914, and 0.925 at 85%, 90%, 95%, and 98% identity, respectively, and outperforming ESM-2, the strongest baseline, by 1.20–2.70 percentage points. With SVM, BacLLM (8B) achieved corresponding mean F1 scores of 0.871, 0.889, 0.900, and 0.919, ranking first at the 85%, 90%, and 98% thresholds. At 95% identity, its performance was only 0.07 percentage points below that of ProTrek. BacLLM (4B) exhibited the same overall advantage, outperforming all baseline models at all four thresholds with both LR and SVM, with mean F1 scores ranging from 0.877 to 0.923 and from 0.873 to 0.924, respectively. These results reproduced the comparative performance pattern observed with the primary 8:2 split, indicating that the relative advantage of BacLLM remained robust under different sequence-identity constraints. Together, these results demonstrate that BacLLM improves protein sequence representations across diverse AMR-related tasks, including ARG identification, gene mobility prediction, resistance class prediction, β-lactamase subtype classification, and resistance mechanism prediction.

### BacLLM captures resistance-associated mutation effects at single-residue resolution

To assess whether BacLLM is sensitive to amino acid substitutions associated with resistance phenotypes, we performed systematic in silico mutagenesis of resistance proteins (Fig. 3a). We introduced single-amino-acid substitutions at each position in the resistance protein sequences to construct full-length mutation landscapes. Based on predictions from the SVM classifier trained for antimicrobial resistance gene (ARG) identification, we calculated the change in predicted ARG probability associated with each amino acid substitution relative to the original sequence (*ΛP* = *P*_mutated_ - *P_initial_*). This score was used as a model-derived proxy to estimate the impact of mutation-associated changes in resistance phenotype.

**Fig. 3.**
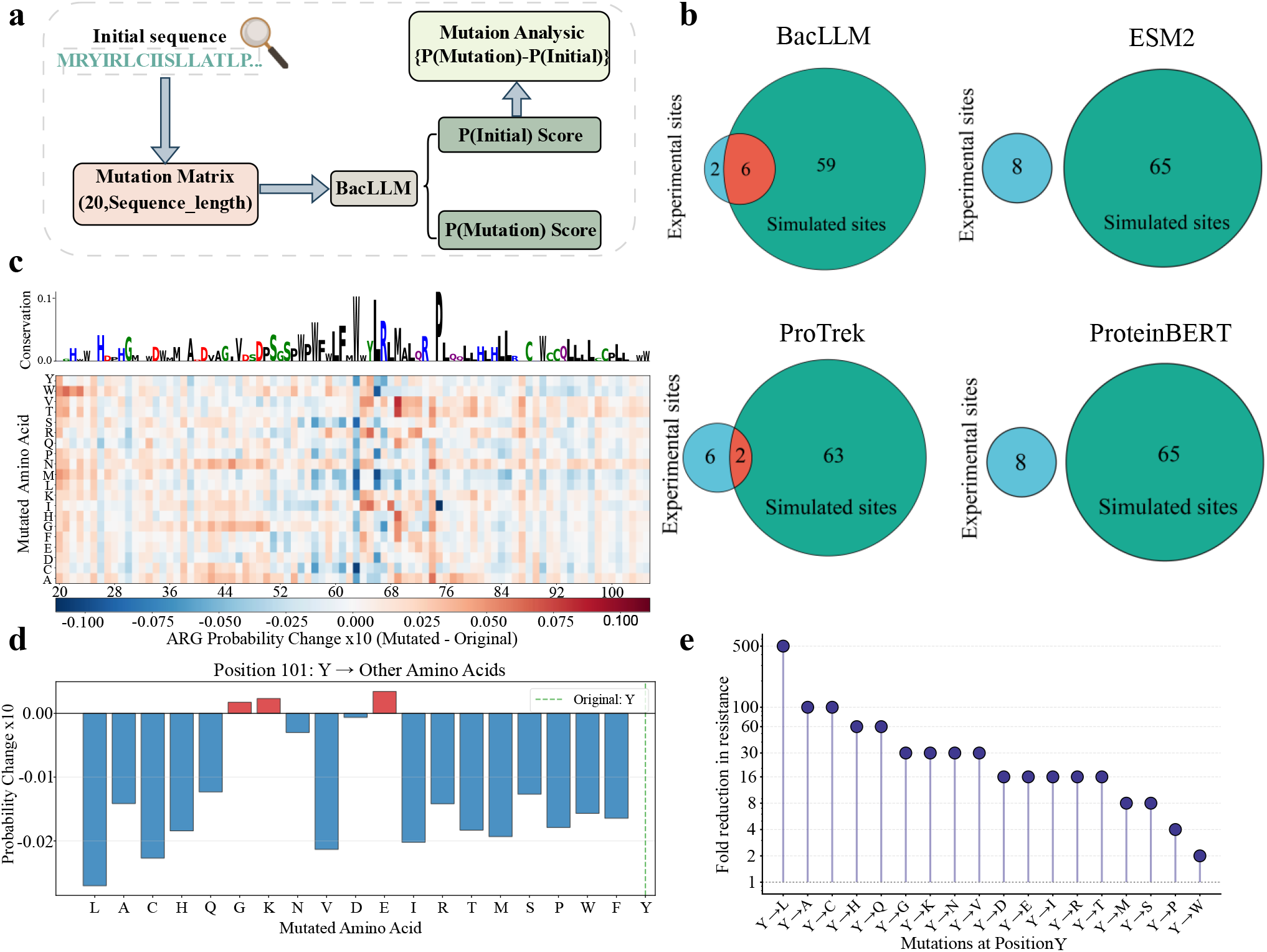
Application of BacLLM in sequence feature analysis. a, quantitative evaluation framework for resistance-associated sequence perturbations using in silico mutagenesis; b, set comparisons of resistance-attenuating mutation sites on the P0AD64 protein predicted by BacLLM, ESM-2, ProTrek, and ProteinBERT against known experimental sites; c, mutational landscape heatmap integrated with sequence conservation, illustrating resistance probability shifts induced by amino acid substitutions; d, detailed mutational profiling at the critical P0AD64 position Y101, showing predicted changes in resistance probability for amino acid substitutions; e, corresponding fold-reductions in resistance reported in the literature for substitutions at the same position.

We first applied this analysis to the well-characterized SHV-1 β-lactamase from *Klebsiella pneumoniae* (UniProt: P0AD64). UniProt records eight experimentally characterized mutation sites in this protein that are associated with reduced resistance or resistance-related functional changes. As shown in Fig. 3b, for each sequence position, we selected the amino acid substitution producing the largest absolute change in predicted resistance probability to represent its dominant mutational effect. We then retained positions whose dominant effects decreased the predicted resistance probability and ranked them by the magnitude of this decrease. Among the top 65 ranked positions, BacLLM (4B) recovered six of the eight experimentally supported resistance-attenuating sites. Under the same evaluation protocol, ESM-2 did not recover any validated sites. Similar analyses of three additional resistance-associated proteins, P52699, P00811, and P02930, produced consistent results (Supplementary Fig. S5). The local mutation landscape further showed that different amino acid substitutions at the same sequence position produced different probability shifts (Fig. 3c). To assess whether these differences were consistent with experimental observations, we examined residue 101 of P0AD64 in greater detail because UniProt provides experimental records for multiple amino acid substitutions at this position. Among the substitutions modelled at this position, BacLLM assigned the largest negative *ΛP* to Y101L, indicating the greatest predicted reduction in resistance probability (Fig. 3d). We then compared this prediction with experimental evidence curated in UniProt, which indicated that substitutions at residue 101 produced different degrees of resistance attenuation, with Y101L causing the largest reduction of approximately 500-fold (Fig. 3e). This experimental observation was consistent with the BacLLM prediction that Y101L produced the largest negative probability shift. Beyond Y101L, BacLLM also broadly captured the experimentally reported variation in resistance attenuation among the other amino acid substitutions at residue 101. Predictions at additional experimentally characterized sites similarly reflected the reported effects of amino acid substitutions on resistance phenotypes (Supplementary Fig. S6). Together, these results support the ability of BacLLM to capture resistance-associated mutation effects at single-residue resolution.

### BacLLM captures taxonomic relationships beyond phenotype-based clinical groupings

We next investigated whether BacLLM embeddings retained signals consistent with known phylogenetic relationships among clinically important bacteria. We averaged the BacLLM-derived protein embeddings within each bacterial strain and used the strain-level embeddings for unsupervised hierarchical clustering of the ESKAPE collection, which comprises six bacterial groups of major clinical relevance to healthcare-associated infections and AMR (Supplementary Table S2). As shown in Fig. 4, the cosine similarity heatmap revealed a structured representation space that broadly reflected known taxonomic relationships among bacterial groups. Strains assigned to the same bacterial group generally formed high-similarity diagonal blocks, indicating strong within-group consistency, with only a small number of *Enterobacter* strains appearing as outliers. At a broader taxonomic level, closely related bacterial groups also showed similar representations. For example, *Klebsiella pneumoniae* and *Enterobacter* spp., both members of Enterobacteriaceae, exhibited high representation similarity. More broadly, Gram-positive and Gram-negative bacteria were largely separated in the representation space, consistent with broad evolutionary differences among these bacterial groups.

**Fig. 4.**
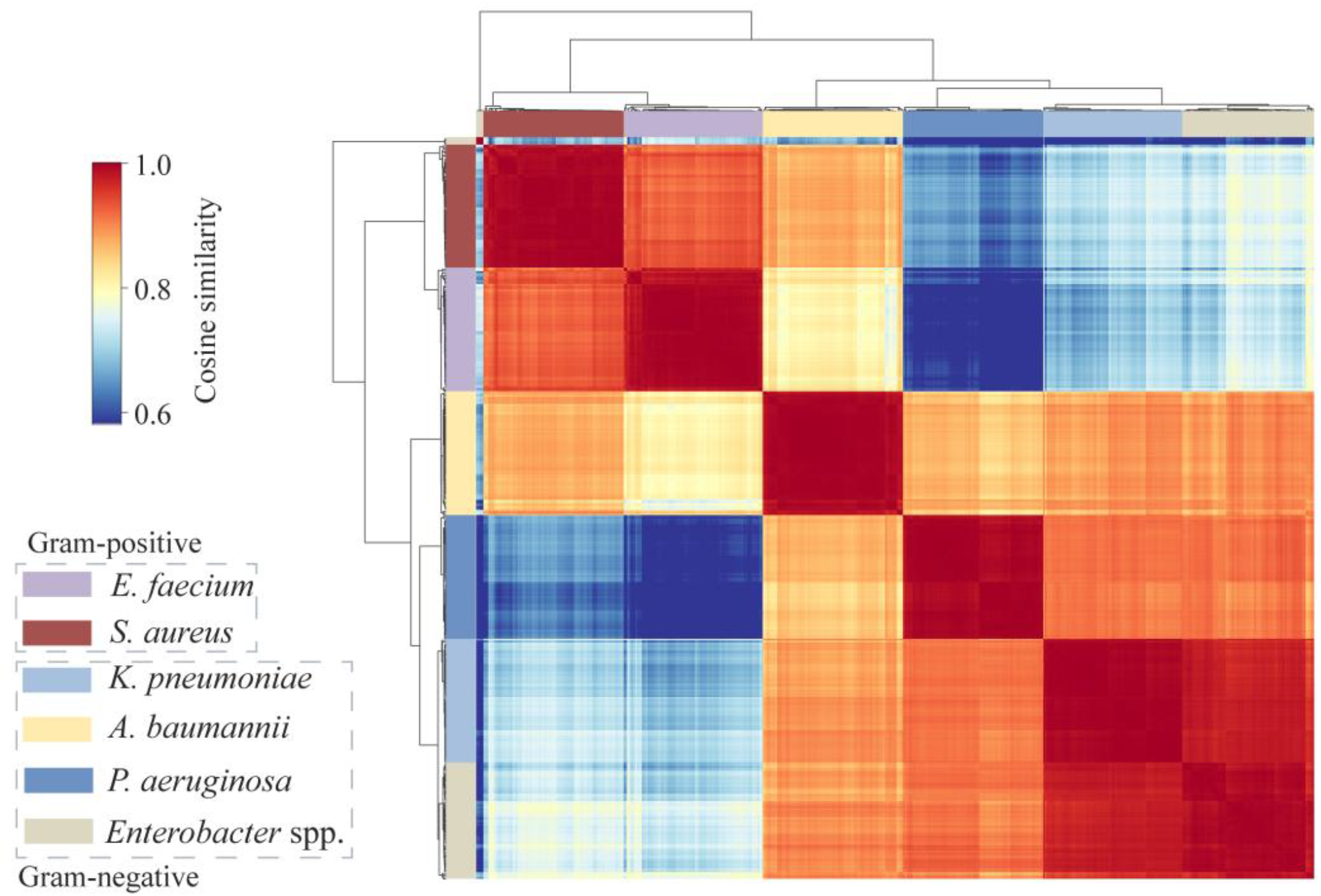
Taxonomic relationships captured by BacLLM representations. Unsupervised phylogenetic clustering of ESKAPE pathogens based on representation similarity.

Notably, BacLLM representations did not simply reflect phenotype-based clinical groupings. In clinical microbiology, *A. baumannii* and *P. aeruginosa* are commonly grouped as major non-fermenting Gram-negative pathogens. However, they formed distinct clusters in the BacLLM representation space. This interpretation is consistent with recent phylogenomic analyses of Pseudomonadales that place Acinetobacter and Pseudomonas in distinct families, Moraxellaceae and Pseudomonadaceae, respectively [35]. Thus, BacLLM representations did not simply follow shared clinical phenotypes but instead exhibited a separation consistent with known phylogenetic relationships.

### BacLLM recovers resistance-associated functional organization in protein networks

Protein association networks provide a means to investigate functional relationships among proteins involved in antimicrobial resistance. To examine whether BacLLM embeddings encode such relationships, we constructed an unsupervised representation-derived protein association network from BacLLM sequence representations. Specifically, we extracted protein embeddings and computed pairwise cosine simi-larities to quantify representation-level associations between proteins. The Leiden algorithm was then applied for community detection, partitioning the network into densely connected communities. For visualization, protein nodes were labeled using their corresponding gene symbols or locus tags.

We focused on TolC, a well-characterized outer membrane channel involved in multidrug efflux and anti-microbial resistance in *Klebsiella pneumoniae* [36]. We identified the representation-derived community containing TolC and extracted a TolC-centered subnetwork (Supplementary Table S5). For visualization, we retained the top 20 ranked proteins and highlighted edges with cosine similarity greater than or equal to 0.65.

As shown in Fig. 5a, the resulting representation-derived network recovered several major components of canonical RND-type efflux systems in Gram-negative bacteria. TolC showed high embedding similarity to the inner-membrane transporter AcrB, while the periplasmic adaptor protein AcrA and other RND efflux system-associated proteins, including AcrD, AcrF, and AcrE, were assigned to the same local community. Given that the AcrAB–TolC complex is a major multidrug efflux system with broad substrate specificity [37], these results suggest that BacLLM representations can recover resistance-associated functional organization using sequence-only inputs at inference, without task-specific protein-interaction or network supervision.

**Fig. 5.**
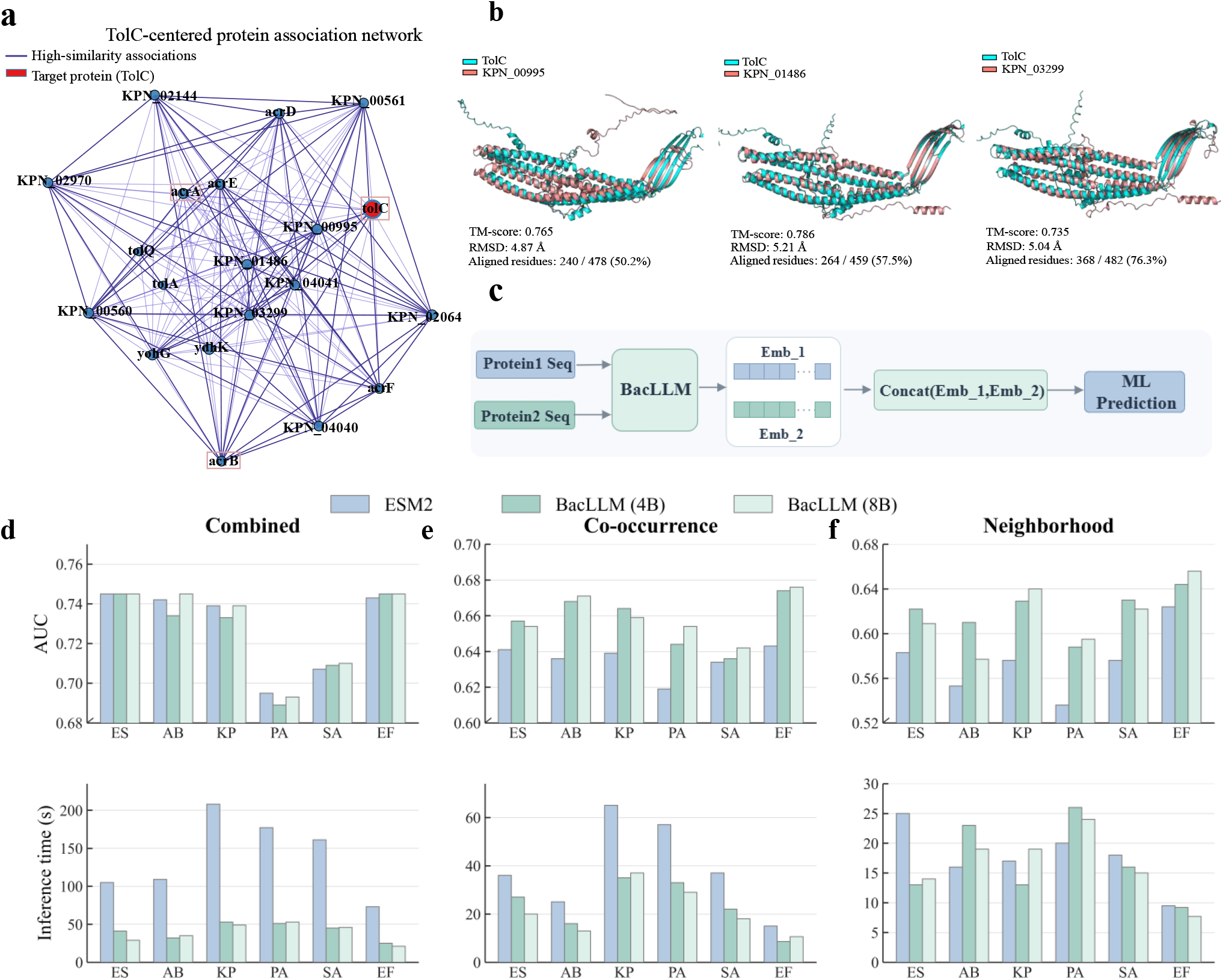
Protein functional association prediction and network analysis. **a**, unsupervised interaction clustering network for the TolC protein based on BacLLM representations; **b**, 3D structural alignments between TolC and three candidate proteins exhibiting high representation similarity within the network**; c**, computational framework for predicting binary protein interactions; **d**–**f**, performance evaluation comparing BacLLM against the baseline ESM-2 in terms of AUC and inference time, evaluated under three distinct interaction types: combined (**d**), co-occurrence (**e**), and neighborhood (**f**).

Having shown that the TolC-centered subnetwork recovered known RND efflux system components, we next investigated whether it could prioritize under-characterized proteins with potential TolC-related proerties. Three proteins, identified by the locus tags KPN_00995, KPN_01486, and KPN_03299, showed high representation similarity to TolC and were retained within the subnetwork. However, these proteins have only limited computational annotations in UniProt-TrEMBL and lack manual curation or experimental validation. KPN_00995 and KPN_01486 are computationally annotated as outer membrane efflux-related proteins, whereas KPN_03299 has only a broad enzyme annotation. This contrast between their close representation-derived association with TolC and limited functional annotation motivated further sequence and structural analyses.

Pairwise sequence alignment revealed low sequence identities of 21.5%, 30.6%, and 23.6% to the reference TolC sequence, respectively. These values fall within or near the sequence-identity range commonly associated with the “twilight zone” of homology-based functional inference [38], where sequence alignment alone may not support reliable functional assignment. We therefore retrieved their AlphaFold-predicted structures and performed structural alignment against TolC. As shown in Fig. 5b, the alignments yielded TM-scores [39] of 0.765, 0.786, and 0.735, with alignment coverages of 50.2%, 57.5%, and 76.3%, respectively, indicating substantial structural similarity to TolC across the aligned regions despite low sequence identity. Together, their placement within the TolC-centered representation network and their structural similarity to TolC support their prioritization as candidate proteins with TolC-like structural features for further functional validation. These findings suggest that BacLLM can complement sequence similarity-based annotation by prioritizing under-characterized proteins for resistance-related functional investigation.

Having demonstrated that BacLLM representations capture resistance-associated functional organization in an unsupervised protein association network, we next evaluated BacLLM in supervised protein functional association prediction using STRING-derived data. As shown in Fig. 5c, due to the large scale of the STRING database, we restricted our evaluation to six clinically important ESKAPE bacteria [23]. We focused on three core types of protein interaction relationships: Combined score, Co-occurrence, and Neighborhood. Given the large dataset size and computational constraints of SVM training, we uniformly used LR as the downstream classifier for comparison. The results are shown in Fig. 5d–f. In the Combined task, BacLLM (8B) achieved performance comparable to state-of-the-art baseline models such as ESM-2. Notably, BacLLM required approximately one-third of the overall computation time of ESM-2, potentially reflecting more efficient optimization of the LR classifier in its representation space. In the Co-occurrence and Neighborhood tasks, BacLLM consistently outperformed ESM-2 across all evaluated bacteria, with absolute AUC improvements of up to 4 and 6 percentage points, respectively, while also demonstrating lower computational cost.

Overall, compared with ESM-2 of similar scale, BacLLM achieved a better trade-off between predictive performance and computational efficiency.

### BacLLM enables accurate phage–host interaction prediction

Phage therapy has attracted increasing attention as a targeted antibacterial strategy for difficult-to-treat bacterial infections associated with AMR [40]. However, phage host ranges are often strain-specific, making it challenging to rapidly identify phages capable of infecting individual bacterial strains [41]. Here, we constructed a dual-branch protein representation pipeline for strain-level phage–host interaction prediction (Fig. 6a), incorporating phage receptor-binding proteins (RBPs) and bacterial surface-associated proteins involved in the initial recognition of bacterial hosts. Detailed sequence processing procedures are provided in Supplementary Note S3.

**Fig. 6.**
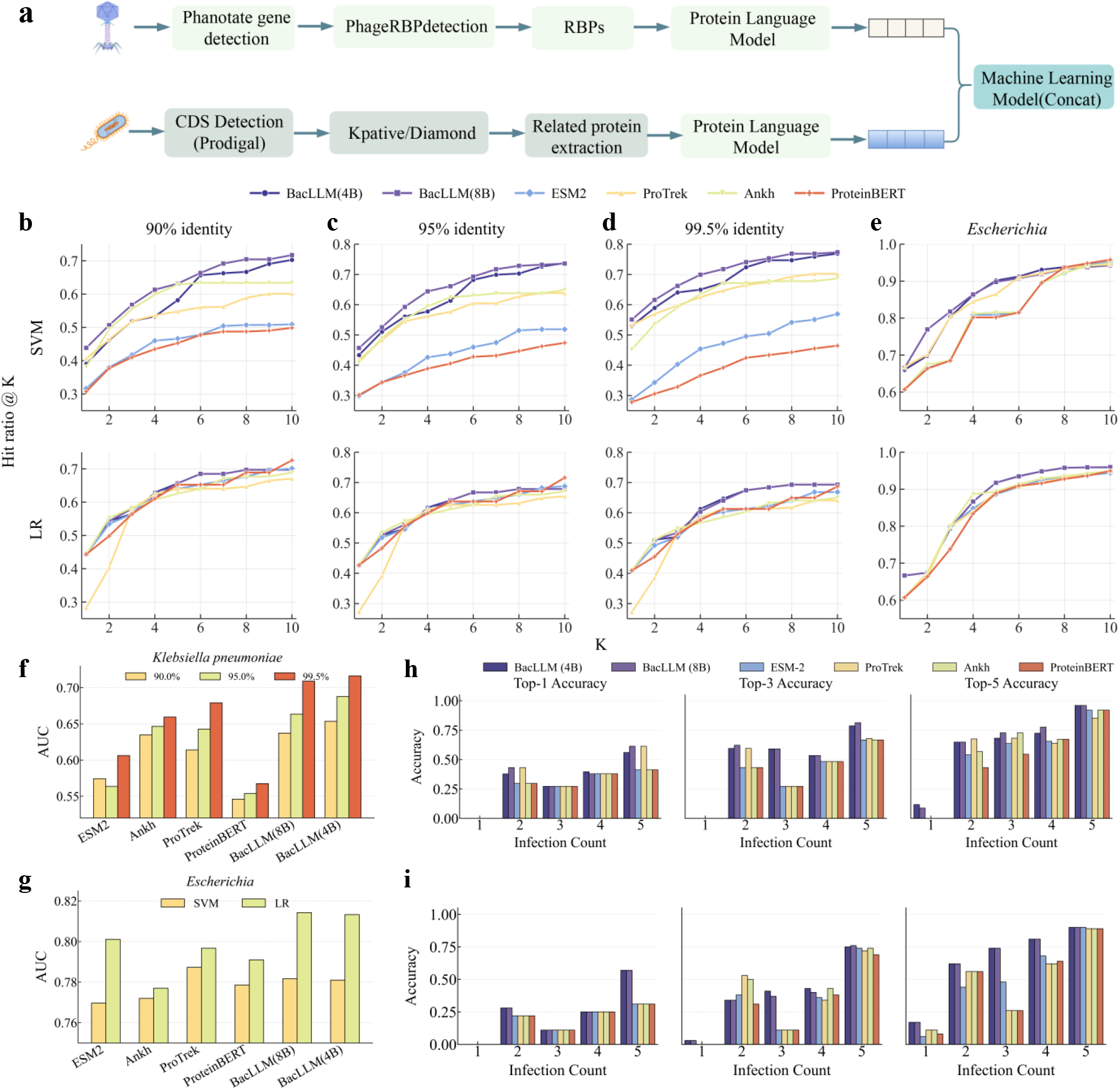
Predictive analysis of specific phage–bacteria interactions. **a**, computational framework for predicting phage–host interactions; **b**–**e**, Hit@K curves utilizing downstream classifiers (top row: SVM; bottom row: LR), evaluated on the Klebsiella pneumoniae dataset at clustering thresholds of 90% (**b**), 95% (**c**), and 99.5% (**d**), and on the *Escherichia* genus dataset (**e**); **f**, **g**, AUC performance comparisons across BacLLM and baseline models for the *Klebsiella pneumoniae* dataset at varying clustering thresholds (**f**) and the *Escherichia* dataset (**g**); **h, i,** sensitivity analysis of Top-K predictive accuracy of BacLLM against base-lines across varying numbers of infecting phages per bacterial host, using SVM (h) and LR (i) as down-stream classifiers.

We evaluated BacLLM on the *Klebsiella pneumoniae* phage–host interaction dataset using the Top-K hit rate as the primary evaluation metric. As shown in Fig. 6b–d, bidirectional best-match clustering was performed at sequence similarity thresholds of 90%, 95%, and 99.5%, followed by leave-one-cluster-out cross-validation (LOCO-CV). With SVM as the classifier, BacLLM (8B) consistently outperformed all baselines across all three clustering thresholds. At the most stringent 90% threshold, BacLLM (8B) achieved a Top-1 hit rate of 43.9%, representing an absolute improvement of approximately 3.2 percentage points over ProTrek, the best-performing baseline. At Top-10, BacLLM (8B) achieved a hit rate of 71.7%, outperforming Ankh, the best-performing baseline, by 8.3 percentage points. Similar trends were observed at other clustering thresholds. At the 95% threshold, BacLLM (8B) outperformed Ankh by 8.7 percentage points at Top-10. At the 99.5% threshold, BacLLM (8B) achieved a Top-10 hit rate of 77.3%, exceeding ProTrek by approximately 7.1 percentage points. When logistic regression (LR) was used as a linear classifier, performance varied across models, and no single model consistently dominated at every Top-K position. Therefore, we quantified overall performance by counting the number of Top-K positions at which each model achieved the highest hit rate. Detailed numerical results are provided in Supplementary Tables S6–S11. The two BacLLM variants demonstrated the strongest overall stability across the entire Top-K range. At the 90% threshold, one or both BacLLM variants achieved or tied for the highest hit rate at seven of the ten evaluated K values. Similar trends were observed at the 95% threshold, where BacLLM achieved or tied for the highest hit rate at six K values. At the 99.5% threshold, BacLLM exhibited a clear advantage, achieving or tying for the highest hit rate at nine of the ten K values. These results indicate that BacLLM maintains competitive matching performance even under the constraints of a linear decision boundary.

To evaluate BacLLM’s crossstrain representation capability, we further analyzed the *Escherichia* dataset. Given its large scale and the substantial computational cost of leave-one-cluster-out cross-validation (LO-CO-CV), we randomly divided the bacterial strains into training and test sets at an 8:2 ratio, ensuring that no strains were shared between the two sets. The experiment was independently repeated 20 times using different random seeds, and the final performance was reported as the average across all runs. As shown in Fig. 6e, with SVM as the classifier, BacLLM exhibited a clear performance advantage across the Top-2 to Top-8 evaluation range. At Top-2, BacLLM (8B) achieved a hit rate of 76.9%, outperforming ProTrek, the best-performing baseline, by 6.8 percentage points. Across the first eight evaluation points, BacLLM (4B) outperformed all baseline models at six positions. Although ProteinBERT and ESM-2 surpassed BacLLM at larger K values (K::9), BacLLM maintained an advantage at smaller K values, where accurate candidate prioritization is particularly important. When LR was used as the classifier, BacLLM achieved or tied for the highest hit rate at eight of the ten evaluated Top-K positions. Notably, at Top-1, both BacLLM (4B) and BacLLM (8B) achieved a hit rate of 66.6%, whereas the best-performing baseline models achieved 60.7%. Detailed numerical results are provided in Supplementary Tables S12–S13.

Overall, Top-K hit rates were generally higher on the *Escherichia* dataset than on the *K. pneumoniae* dataset. This difference may partly reflect the denser host-range matrix of the Escherichia dataset, which contains a higher proportion of phages associated with multiple bacterial hosts and may therefore provide richer interaction information for model learning. However, the two datasets were evaluated using different data-splitting protocols, which should also be considered when interpreting their performance differences. Collectively, the results across these two clinically important bacterial groups demonstrate the potential of BacLLM for strain-level phage–host matching and suggest that improved candidate prioritization could reduce the number of phages requiring downstream experimental validation, highlighting its potential utili-ty for large-scale phage–host screening.

To further evaluate pairwise classification performance, we used AUC as a complementary evaluation metric. As shown in Fig. 6f, performance on the *K. pneumoniae* dataset was assessed using SVM, which demonstrated stronger overall performance than LR in the preceding Top-K evaluation across the three sequence similarity thresholds of 90%, 95%, and 99.5%. As expected, prediction difficulty increased as the similarity threshold decreased. Nevertheless, BacLLM remained robust and consistently outperformed all baseline models across thresholds. At the 99.5% threshold, BacLLM (4B) and BacLLM (8B) achieved AUC values of 0.72 and 0.71, respectively, each representing an absolute improvement of approximately 10 percentage points over ESM-2. The results on the *Escherichia* dataset are shown in Fig. 6g. When LR was used as the classifier, BacLLM (8B) achieved the highest AUC, reaching 0.81. When SVM was used, all models achieved highly similar performance, with ProTrek showing a slight advantage.

To further assess the model’s ability to capture highly specific phage–host interactions, we focused on matching tasks involving low-susceptibility bacteria. From a biological perspective, lower host susceptibil-ity is often associated with more stringent receptor–ligand recognition requirements, resulting in sparser positive interactions and a substantially more challenging prediction task. We therefore selected bacterial strains that could be infected by only one to five phages and systematically evaluated Top-1, Top-3, and Top-5 matching hit rates across different susceptibility levels. As shown in Fig. 6h,i, BacLLM remained robust under both SVM (Fig. 6h) and LR (Fig. 6i), particularly for hosts susceptible to very few phages. In the most stringent setting, where each host was susceptible to only one phage, BacLLM was the only representation that enabled successful retrieval: the correct phage appeared within the Top-5 candidates using SVM and within the Top-3 candidates using LR, whereas all baseline models failed. The advantage generally persisted as host susceptibility increased. With LR, BacLLM exceeded the best-performing baseline, ProTrek, by approximately 6 percentage points in Top-5 hit rate for hosts susceptible to two phages; when susceptibility increased to five phages, its Top-1 hit rate reached 0.57, compared with approximately 0.31 for ESM-2, ProTrek, and Ankh. With SVM, BacLLM (8B) achieved a Top-1 hit rate of 0.61 for hosts susceptible to five phages, tying ProTrek and outperforming the other baselines (approximately 0.41), while the BacLLM variants achieved the highest Top-5 hit rate of 0.96. Together, these results indicate that BacLLM captures sparse phage–host interaction signals robustly across different downstream classifiers.

In summary, under the highly challenging low-susceptibility setting, BacLLM consistently outperformed existing protein language models in modeling phage–host specificity, highlighting its ability to support phage therapy candidate prioritization by capturing highly specific phage–host interaction signals.

### Ablation Study

To evaluate the contributions of the sequence enhancement module G(0) and the text semantic enhancement module, we conducted ablation experiments on ARGNet-DB under both the Qwen3-4B and Qwen3-8B training settings. The objective was to assess how different module configurations affected the quality of BacLLM representations. As summarized in Supplementary Tables S14–S15, jointly incorporating both modules consistently improved feature separability, with the largest gains observed when SVM was used as the downstream classifier. Compared with the baseline model without enhancement modules, the dual-enhancement configuration improved the F1 score by approximately 4 percentage points in both the 4B and 8B settings. In contrast, introducing either module alone did not yield consistent gains and, in some settings, slightly reduced performance relative to the baseline. These results suggest that enhancing only a single modality may disturb the alignment between sequence and text representations, leading to suboptimal feature distributions. When LR was used as the downstream classifier, performance differences among different module combinations became considerably smaller. This may be because LR had already approached its performance ceiling on this dataset. Nevertheless, the dual-enhancement configuration consistently achieved the best overall performance with LR, further supporting the effectiveness of jointly enhanced representations across different classification models.

We next investigated the effect of scaling the sequence encoder from ESM-2 33M to ESM-2 150M (Supplementary Fig. S7a, b). On both the HMD-ARG and ARGNet datasets, ESM-2 and BacLLM generally benefited from increasing the sequence encoder size when evaluated with LR, with BacLLM consistently outperforming the corresponding ESM-2 baselines. Under SVM, however, ESM-2 performance declined as the encoder size increased, whereas BacLLM maintained or improved its performance. This may reflect the increased difficulty of learning decision boundaries from higher-dimensional frozen embeddings, as the feature dimension increased from 480 in ESM-2 33M to 640 in ESM-2 150M. In contrast, BacLLM was less affected by this dimensional expansion and continued to benefit from the larger sequence encoder. In particular, on HMD-ARG with SVM, BacLLM 150M trained with Qwen3-8B improved the F1 score from 0.902 to 0.944 relative to its 33M counterpart, a gain of 4.2 percentage points. These results suggest that domain-adaptive semantic supervision helps BacLLM maintain stable and discriminative representations as encoder size increases.

We further evaluated model robustness in the *Klebsiella pneumoniae* phage–host interaction task at the 99.5% sequence similarity threshold using SVM (Supplementary Fig. S7c, d). Because this task concatenates phage and bacterial protein embeddings, the input dimensionality increased substantially to 1,280. Under AUC evaluation, the effect of increased dimensionality differed among the models. The AUC of ESM-2 decreased from 0.6062 to 0.5143, a decline of 9.2 percentage points. In contrast, BacLLM (4B) increased from 0.7163 to 0.7263, whereas BacLLM (8B) decreased from 0.7163 to 0.6823, corresponding to changes of +1.0 and −3.4 percentage points, respectively. In fine-grained Hit@K retrieval evaluations, BacLLM also maintained a clear advantage over ESM-2 despite the increased feature dimensionality.

In summary, these results indicate that, in our frozen-encoder evaluation setting, increasing sequence encoder size did not always lead to proportional downstream performance gains. The accompanying increase in feature dimensionality may make downstream classification more difficult, particularly when embeddings are concatenated across phage and bacterial features. Compared with ESM-2, BacLLM showed greater robustness to this dimensional expansion and maintained stable, discriminative representations under high-dimensional feature concatenation.

### Conclusion

BacLLM is an AMR-focused protein representation model adapted to WHO priority bacterial pathogens. It combines sequence enhancement, semantic enhancement of UniProt annotations, and cross-modal contrastive learning to transfer annotation-derived biomedical knowledge into an ESM-2 sequence encoder. During training, protein sequence embeddings are aligned with semantic representations of curated annotations encoded by a large language model. After training, the text branch is removed, and only the sequence encoder is retained. This design separates annotation-guided knowledge transfer during training from input requirements at inference, enabling BacLLM to provide annotation-informed representations from amino acid sequences alone.

We first evaluated whether these representations support a progressive workflow for AMR sequence analysis, extending from ARG identification to resistance function and dissemination potential. Across five biological prediction tasks and six primary benchmark settings, BacLLM generally provided more informative representations of AMR-associated protein sequences than the baseline protein language models under a standardized frozen-encoder evaluation protocol. The ablation results further demonstrated that sequence enhancement and annotation semantic enhancement produced the greatest benefit when applied jointly, whereas either module alone yielded less consistent improvements. These findings highlight the importance of coordinated sequence and annotation enhancement within the cross-modal alignment framework for learning informative AMR-related protein representations.

Beyond conventional AMR-related classification tasks, BacLLM captured biologically meaningful organization at complementary levels. Using changes in ARG classifier-predicted probabilities as a model-derived proxy for resistance-associated effects, the single-residue analysis identified mutational patterns consistent with experimentally reported resistance-attenuating sites. This result indicates that BacLLM representations are sensitive to local sequence perturbations relevant to resistance. When protein representations were aggregated by bacterial strain, BacLLM also preserved broad taxonomic relationships among ESKAPE bacteria, including distinctions not necessarily reflected by phenotype-based clinical groupings. This suggests that AMR-focused adaptation retained general biological organization in the representation space. At the protein-network level, a TolC-centered representation-derived protein association network recovered known components of RND-type efflux systems without task-specific protein-interaction or network supervision. Within this network, BacLLM prioritized under-characterized proteins with low sequence identity to TolC but substantial predicted structural similarity across the aligned regions. The convergence of representation-level associations and structural evidence supports the prioritization of these proteins for further functional investigation.

Having established its utility for AMR characterization, we next evaluated whether BacLLM could support an intervention-oriented application. Using representations of phage receptor-binding proteins and bacterial receptor-associated protein sets, BacLLM improved strain-level phage–host matching across the *Klebsiella pneumoniae* and *Escherichia* datasets. The model also maintained an advantage in low-susceptibility settings, where positive phage–host interactions were sparse and candidate ranking was more challenging. These findings extend the application of BacLLM beyond resistance sequence characterization to phage therapy candidate prioritization. In practice, improved computational ranking could narrow the set of candidate phages advanced to downstream experimental testing, potentially increasing the efficiency of large-scale phage–host screening.

Collectively, BacLLM provides a clinically relevant computational protein representation framework that connects sequence-based AMR surveillance and resistance mechanism characterization with intervention-oriented candidate prioritization. Because it requires only amino acid sequences during inference, BacLLM could serve as a reusable computational layer within future clinical microbiology workflows, supporting the prioritization of resistance-associated sequences, functionally relevant proteins, and candidate phages for confirmatory testing. Several limitations should nevertheless be considered. First, training and evaluation were based on retrospective public datasets; temporally independent and clinically derived datasets will be required to establish generalizability and reduce dataset-specific effects. Second, the prioritized TolC-like candidates remain computational predictions and require direct functional validation, although the mutation and phage–host analyses were evaluated using experimentally derived evidence. Third, alt-hough BacLLM is trained through cross-modal alignment, its downstream analyses currently rely on protein sequence information alone and do not incorporate complementary omics or clinical data. Future work could integrate these additional data sources to support a more comprehensive characterization of AMR mechanisms across biological and clinical conditions.

## Code Availability

The source code and implementation of BacLLM are publicly available on GitHub at https://github.com/oooo111/BacLLM_model/tree/main.

## Data Availability

Protein sequences and functional annotations used for domain-adaptive training were obtained from UniProt (https://www.uniprot.org/). Additional bacterial protein sequences used for benchmark construction were retrieved from the NCBI database (https://www.ncbi.nlm.nih.gov/protein/).

The antimicrobial resistance datasets used in this study were derived from publicly available resources, including HMD-ARG (http://www.cbrc.kaust.edu.sa/HMDARG/), ARGNet-DB (https://github.com/id-bioinfo/ARGnet), and the Beta-Lactamase Database (BLDB; http://bldb.eu/). Protein functional association data were obtained from STRING (https://string-db.org/cgi/download).

The *Klebsiella pneumoniae* phage–host interaction dataset was obtained from PhageHostLearn and is available through Zenodo (https://doi.org/10.5281/zenodo.11061100). The Escherichia phage–host interaction dataset is publicly available through Zenodo (https://doi.org/10.5281/zenodo.13831957).

## Author Contributions

Rongye Ye conceived the research idea, designed the experiments, and wrote the original draft of the manuscript. Lun Li contributed to algorithm optimization, participated in discussions of key methodological aspects, and reviewed and edited the manuscript. Zongyi Geng proposed the incorporation of an enhancement module and provided valuable insights into the methodological design. Shuhui Song contributed to the conceptual development of the study, established the overall research framework, critically revised the manuscript, and supervised and coordinated the project.

## Supplementary Figure and Table Legends

**Supplementary Table S1.** WHO Bacterial Priority Pathogens List.

**Supplementary Table S2.** Number of ESKAPE samples used in the PPI prediction.

**Supplementary Table S3–S4.** Experiments on the antimicrobial resistance prediction task.

**Supplementary Table S5.** Pairwise cosine similarities between proteins in the TolC-centered protein association network.

**Supplementary Table S6.** In the *Klebsiella pneumoniae* dataset, clustering was performed using a 90% sequence similarity threshold, and LR was employed as the classifier in downstream tasks.

**Supplementary Table S7:** In the *Klebsiella pneumoniae* dataset, clustering was performed using a 90% sequence similarity threshold, and SVM was employed as the classifier in downstream tasks.

**Supplementary Table S8:** In the *Klebsiella pneumoniae* dataset, clustering was performed using a 95% sequence similarity threshold, and LR was employed as the classifier in downstream tasks.

**Supplementary Table S9:** In the *Klebsiella pneumoniae* dataset, clustering was performed using a 95% sequence similarity threshold, and SVM was employed as the classifier in downstream tasks.

**Supplementary Table S10:** In the *Klebsiella pneumoniae* dataset, clustering was performed using a 99.5% sequence similarity threshold, and LR was employed as the classifier in downstream tasks.

**Supplementary Table S11:** In the *Klebsiella pneumoniae* dataset, clustering was performed using a 99.5% sequence similarity threshold, and SVM was employed as the classifier in downstream tasks.

**Supplementary Table S12:** In the *Escherichia* dataset, experiments were repeated using 20 random seeds, and the results were averaged across runs. LR was employed as the classifier in downstream tasks.

**Supplementary Table S13:** In the *Escherichia* dataset, experiments were repeated using 20 random seeds, and the results were averaged across runs. SVM was employed as the classifier in downstream tasks.

**Supplementary Table S14:** Parameter ablation analysis of the BacLLM (4B) model.

**Supplementary Table S15:** Parameter ablation analysis of the BacLLM (8B) model.

**Supplementary Fig. S1** | Textual Semantic Enhancement Module. Textual semantic enhancement was performed using the MedLLM-Llama3-8B model. Through carefully designed prompt engineering, these fragmented attribute features were reconstructed into logically coherent and semantically enriched natural language descriptions.

**Supplementary Fig. S2** | Class distributions of antimicrobial resistance datasets used in this study. a, distribution of resistance categories in the HMD-ARG dataset; b, distribution of resistance categories in the ARGNet dataset; c, distribution of resistance mechanisms in the HMD-ARG dataset; d, distribution of β-lactamase subclasses in the BLDB database.

**Supplementary Fig. S3** | Visualization of phage–host interaction relationships. a, illustrative phage–host interaction network derived from a subset of *Klebsiella pneumoniae* data. b, illustrative phage–host interaction network derived from a subset of *Escherichia* data.

**Supplementary Fig. S4** | Performance efficiency comparison of protein language models across four sequence-identity thresholds using MMseqs2-guided five-fold cross-validation.

**Supplementary Fig. S5** | Extended evaluation of mutation effect prediction for resistance-associated proteins. a, overlap between experimentally validated functional mutation sites and the top-ranked mutation sites predicted by the model for protein P52699. b, corresponding results for protein P00811. c, corresponding results for protein P02930.

**Supplementary Fig. S6** | Additional analysis of mutation sites in P0AD64. We further examined four additional mutation sites that were independently identified by both experimental studies and model predictions. a, residue 66; b, residue 270; c, residue 126; d, residue 230.

**Supplementary Fig. S7** | Extended scaling analysis of BacLLM. a, Resistance category prediction performance on the HMD-ARG dataset. b, Resistance category prediction performance on the ARGNet dataset. c, AUC performance on the Klebsiella pneumoniae phage–host interaction dataset at a 99.5% sequence similarity threshold. d, Hit@K performance on the same dataset. SVM was used as the downstream classifier in c and d.

