## Supplementary information for "Language-guided domain adaptation improves protein sequence representations for antimicrobial resistance analysis"

**The file include:**

**Supplementary Notes S1-S3**

**Supplementary Figures S1-S7**

**Supplementary Tables S1-S15**

Supplementary Note S1

To quantify the similarity of K-locus regions across different bacterial strains and perform clustering analysis, we employed a similarity assessment and hierarchical clustering framework based on protein sequence multisets. Specifically, each strain $X_{i}$ was defined as a multiset containing $n_{i}$ proteins encoded within its K-locus region:

$$X_{i}= \{p_{\left\{ i,1 \right\}}, p_{\left\{ i,2 \right\}}, \cdots, p_{\left\{ i,n_{i} \right\}}\}$$

where $p_{i,k}$ denotes the kth K-locus protein sequence in strain $X_{i}$, and $n_{i}$ represents the total number of K-locus proteins contained in that strain. To quantify the similarity between two individual protein sequences $a$ and $b$, we introduced a normalized identity function $f_{id}\left( a,b \right)$ based on the Levenshtein edit distance:

$$f_{id}\left( a,b \right)=1-\frac{d_{lev}\left( a,b \right)}{\max\left( \left| a \right|,\left| b \right| \right)}$$

In this formulation, $d_{lev}\left( a,b \right)$ denotes the Levenshtein edit distance between sequences $a$ and $b$, defined as the minimum number of single-character edit operations required to transform one sequence into the other. $\left| a \right|$ and $\left| b \right|$ represent the lengths of sequences $a$ and $b$, respectively. This normalization ensures that the value of $f_{id}$ ranges from 0 to 1. Building upon pairwise sequence similarity, we next quantified the overall similarity $Sim\left( X_{A},X_{B} \right)$between two strains, $X_{A}$ and $X_{B}$, using a bidirectional best-match strategy combined with a length penalty term to account for differences in the numbers of K-locus proteins:

$$Sim\left( X_{A},X_{B} \right)=\frac{\sum_{a\in X_{A}} \max_{b\in X_{B}} f_{id}\left( a,b \right)+\sum_{b\in X_{B}} \max_{a\in X_{A}} f_{id}\left( b,a \right)}{2\cdot\max\left( \left| X_{A} \right|,\left| X_{B} \right| \right)}$$

In the numerator, the first term sums the highest similarity score obtained for each protein in $X_{A}$ against all proteins in $X_{B}$, whereas the second term performs the reciprocal calculation for proteins in $X_{B}$ against those in $X_{A}$. This bidirectional matching scheme ensures a symmetric and balanced assessment of strain similarity. The denominator, $2\max\left( \left| X_{A} \right|,\left| X_{B} \right| \right)$, serves as a penalty term that mitigates the influence of substantial differences in the numbers of K-locus proteins carried by the two strains. Based on the strain similarity scores defined above, we constructed a pairwise distance matrix, in which the distance $D_{ij}$ between any two strains $X_{i}$ and $X_{j}$ was defined as:

$$D_{ij}=1-Sim\left( X_{i},X_{j} \right)$$

Hierarchical clustering was subsequently performed using the resulting distance matrix. The clustering procedure employed the complete-linkage criterion, in which the distance $d\left( C_{u},C_{v} \right)$ between two clusters $C_{u}$ and $C_{v}$ was defined by the maximum pairwise distance between any two members belonging to the respective clusters:

$$d\left( C_{u},C_{v} \right)=\max_{i\in C_{u},j\in C_{v}} D_{ij}$$

Finally, clusters were determined by cutting the resulting dendrogram at a predefined threshold $\tau$.

Supplementary Note S2

TM-score is a standard metric for evaluating the global topological similarity between two protein structures. Its mathematical formulation is defined as follows:

$$\text{TM-score}=\max\left[ \frac{1}{L_{\text{target}}}\sum_{i=1}^{L_{\text{ali}}} \frac{1}{1+\left( \frac{d_{i}}{d_{0}\left( L_{\text{target}} \right)} \right)^{2}} \right]$$

$$d_{0}\left( L_{\text{target}} \right)=1.24\sqrt[3]{L_{\text{target}}-15}-1.8$$

In this formulation, $d_{0}\left( L_{\text{target}} \right)$ is a length-dependent scaling factor that dynamically adjusts according to protein size, thereby providing a consistent evaluation criterion across proteins of different lengths. $L_{target}$ denotes the total amino acid sequence length of the target protein, whereas $L_{ali}$ represents the number of aligned residue pairs between the two structures after structural superposition. $d_{i}$is the Euclidean distance between the $i$th pair of aligned residues following optimal structural alignment.

Supplementary Note S3

In the phage branch, open reading frames were identified from phage genomes using PHANOTATE [1], and RBP sequences were subsequently detected using PhageRBPdetection [2]. The identified RBPs were then encoded using pretrained protein language models. In the bacterial branch, receptor-associated protein features were selected according to the biological characteristics of each dataset. For Klebsiella pneumoniae, capsule synthesis loci (K-loci) were identified using Kaptive [3], and the corresponding K-locus proteins were used as sequence-derived proxies for capsular polysaccharide composition, a major determinant of phage adsorption and host specificity for many Klebsiella phages [4]. For Escherichia, previous work identified variation in bacterial surface polysaccharides, including LPS outer-core type, O-antigen serotype, and capsule type, as a major correlate of strain-level phage susceptibility [5]. These surface structures can serve as primary receptors or modulate access to other receptors, thereby influencing phage adsorption. Accordingly, host representations included outer membrane proteins, some of which can act as direct receptors for specific phages [6], together with proteins involved in LPS and capsule biosynthesis and export. The latter proteins were used as sequence-derived proxies for strain-level variation in surface polysaccharide composition. To identify these proteins, we constructed a reference database using receptor-associated proteins from the reference strain Escherichia coli K-12. Protein-coding genes were predicted using Prodigal [7], and the resulting protein sequences were searched against this reference database using DIAMOND [8] to identify receptor-associated homologues.

Supplementary materials

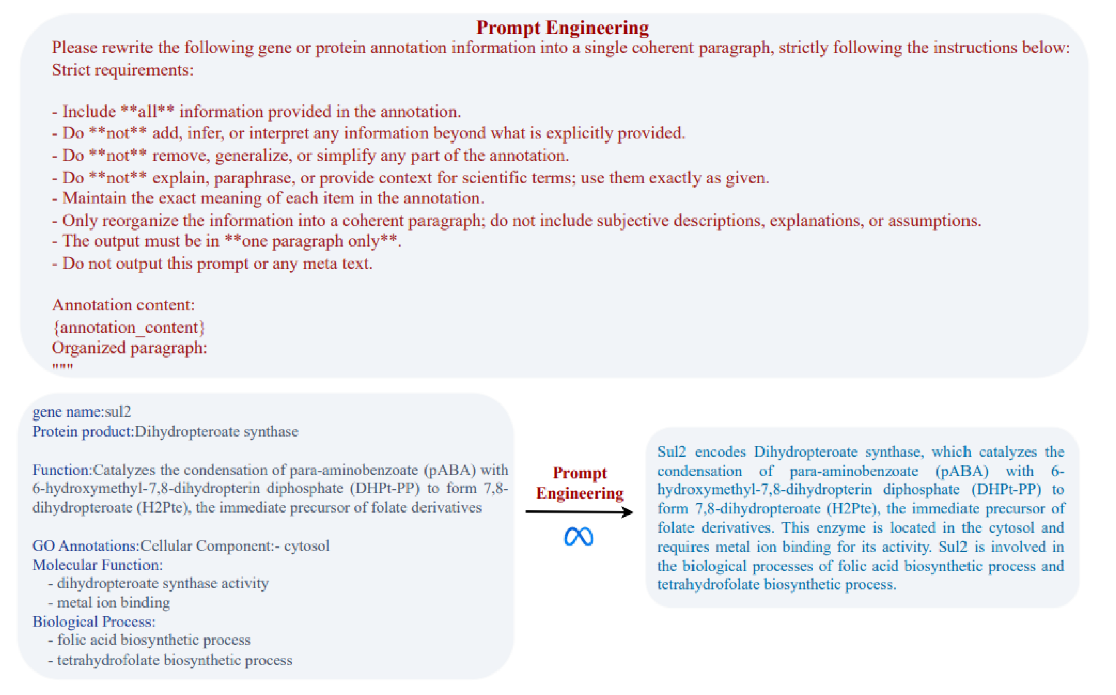

**Note:** Text semantic enhancement was performed using the **MedLLM-Llama3-8B** model, which is specifically trained for the biomedical domain.

**Supplementary Fig. S1:** Text semantic enhancement module

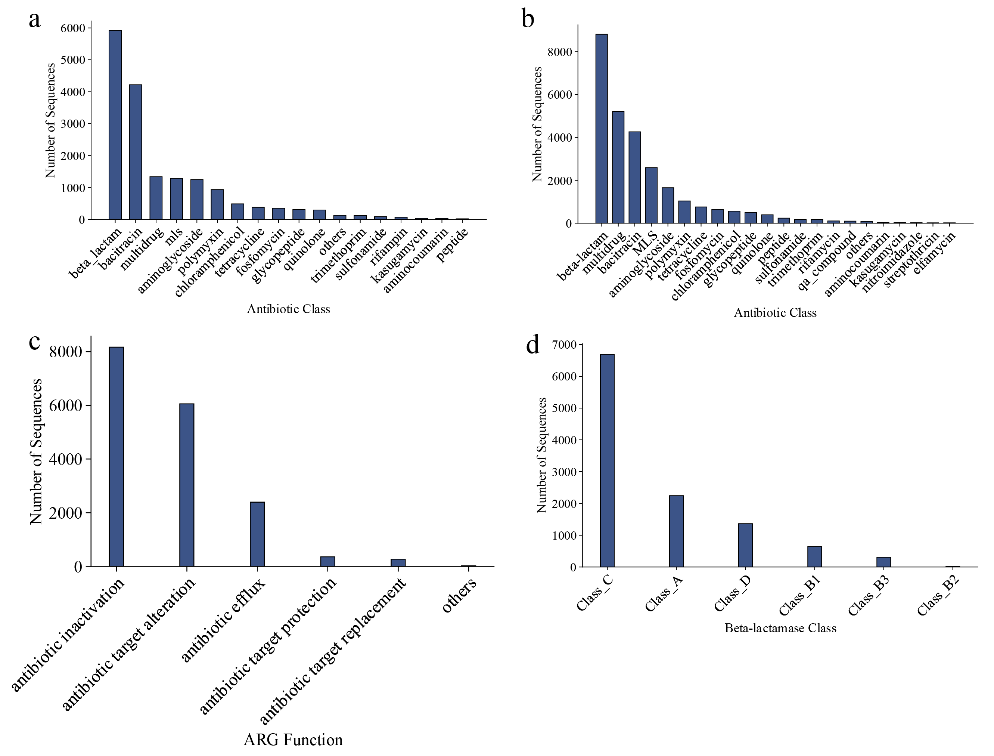

**Supplementary Fig. S2:** Class distributions of antimicrobial resistance datasets used in this study

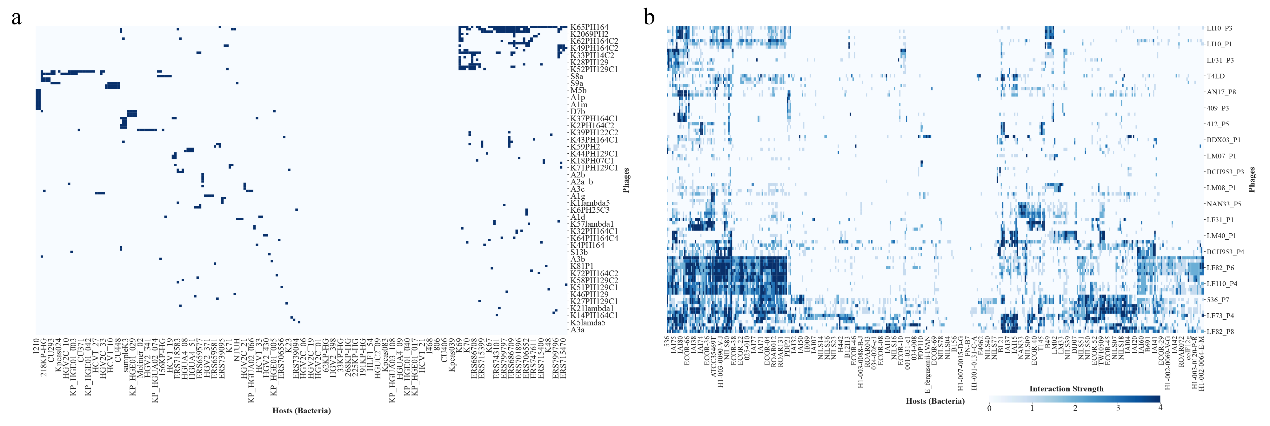

**Supplementary Fig. S3:** Visualization of a subset of phage–host interactions in the *Klebsiella pneumoniae* dataset (left) and the *Escherichia* genus dataset (right)

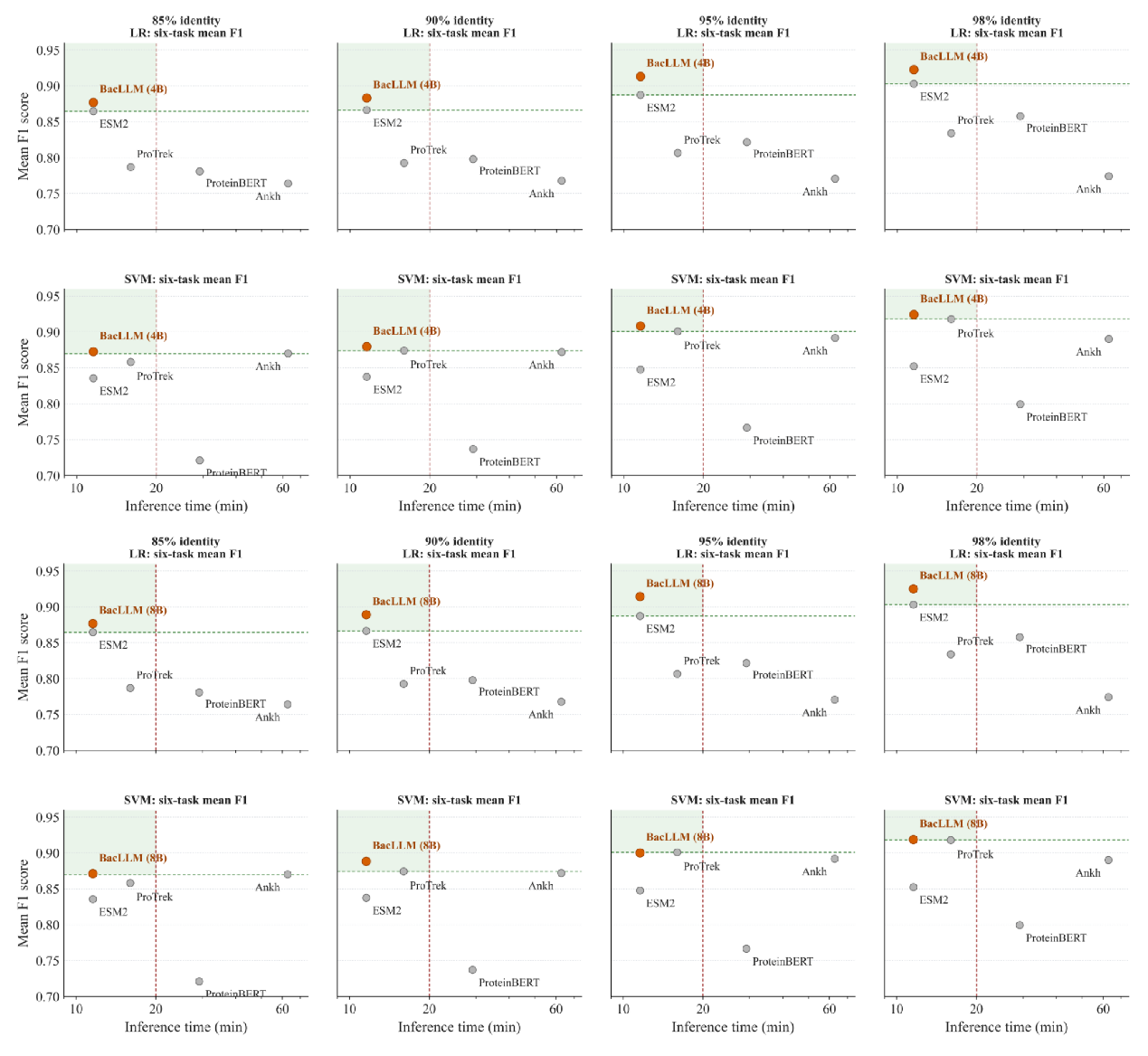

**Supplementary Fig. S4:** Performance efficiency comparison of protein language models across four sequence-identity thresholds using MMseqs2-guided five-fold cross-validation.

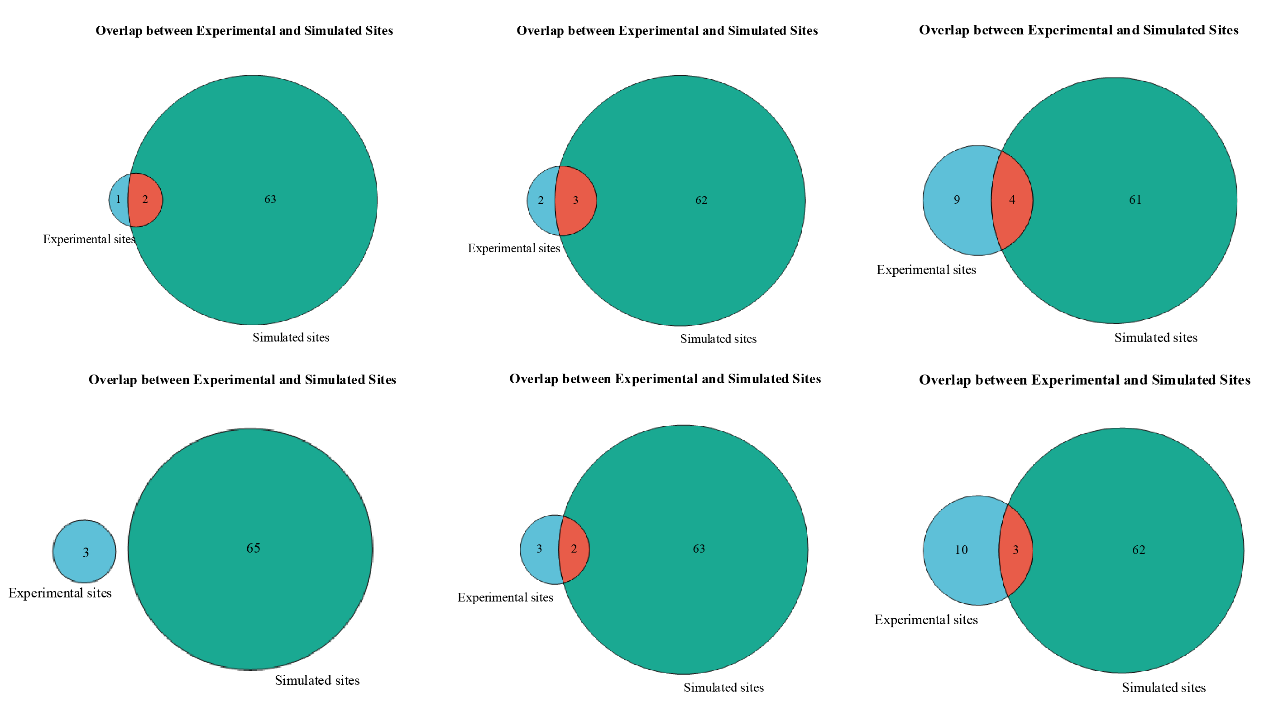

**Supplementary Fig. S5:** Extended evaluation of mutation effect prediction for resistance-associated proteins P52699, P00811, and P02930 (left to right), with results from BacLLM (top) and ESM-2 (bottom).

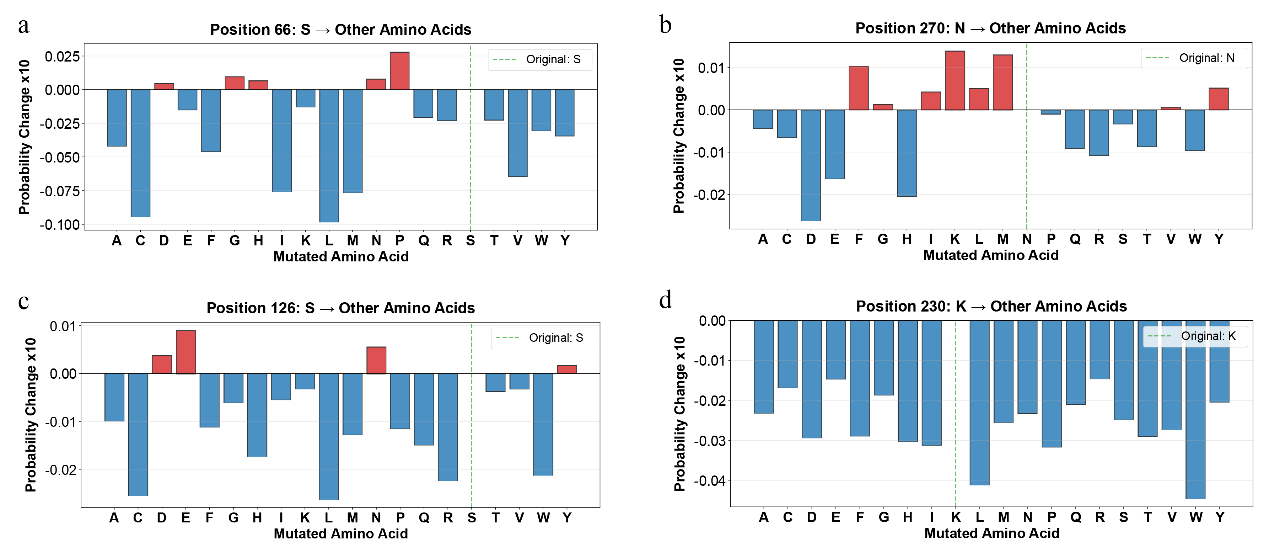

**Supplementary Fig. S6:** Additional analysis of mutation sites in P0AD64.

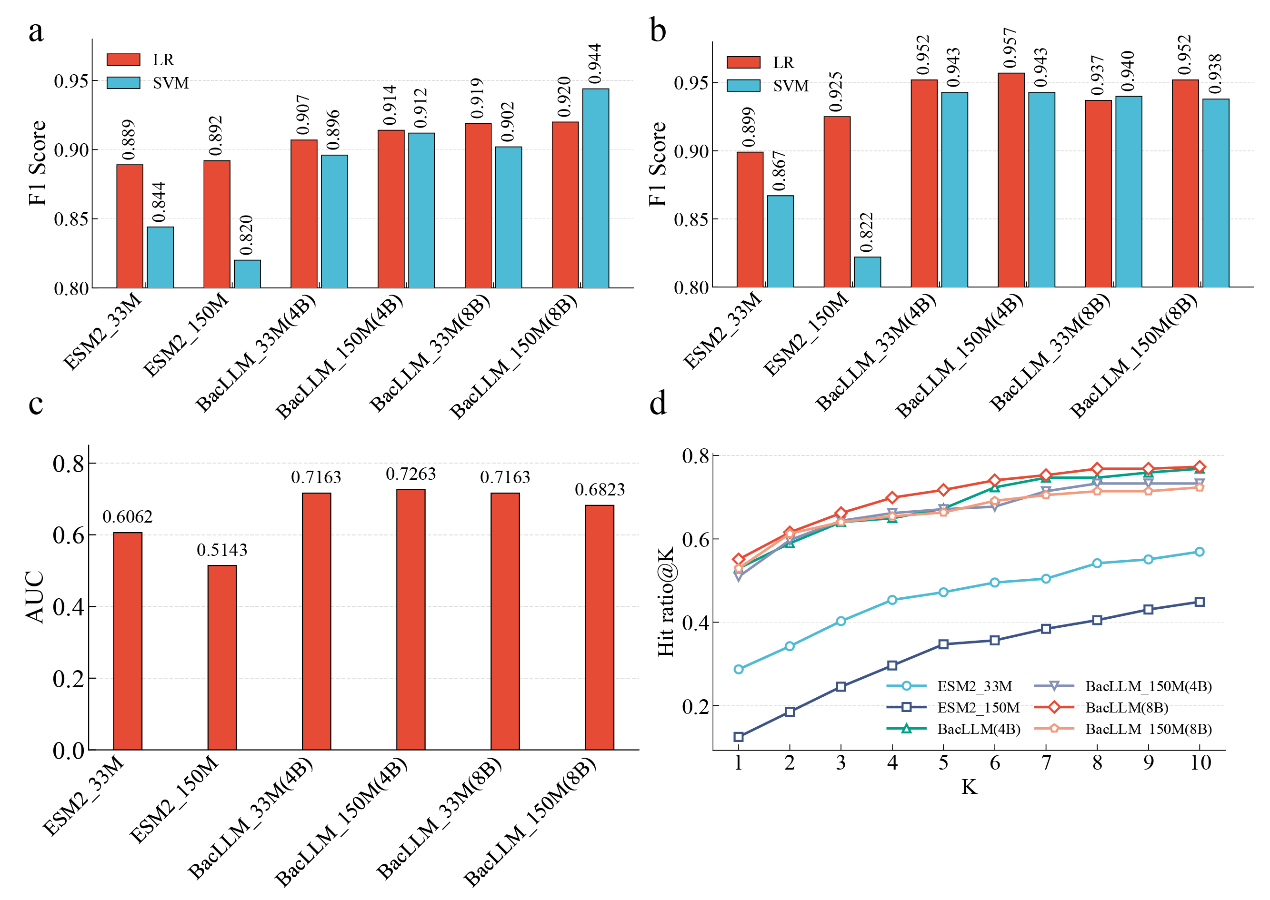

**Supplementary Fig. S7:** Extended ablation and scaling analysis of BacLLM.

**Supplementary Table S1:** WHO Bacterial Priority Pathogens List.

| **Acinetobacter baumannii** | **Enterobacterales** | **Mycobacterium tuberculosis** | **Salmonella Typhi** | **Shigella spp** |
| --- | --- | --- | --- | --- |
| **Enterococcus faecium** | **Pseudomonas aeruginosa** | **Non-typhoidal Salmonella** | **Neisseria gonorrhoeae** | **Staphylococcus aureus** |
| **Group A Streptococci** | **Streptococcus pneumoniae** | **Haemophilus influenzae** | **Group B Streptococci** |  |

**Note:** Red indicates pathogens classified in the WHO Critical group, purple represents the High group, and blue denotes the medium group.

**Supplementary Table S2:** Sample sizes used for PPI prediction across ESKAPE bacteria.

| Bacteria | Acinetobacter baumannii | Enterococcus faecium | Enterobacter species | Klebsiella pneumoniae | Pseudomonas aeruginosa | Staphylococcus aureus |
| --- | --- | --- | --- | --- | --- | --- |
| Numbers | 761,752 | 453,516 | 947,562 | 1,412,338 | 1,827,886 | 862,170 |

**Note:** ESKAPE pathogens are among the most clinically important bacteria associated with hospital-acquired infections and antimicrobial resistance, representing a major threat in the current AMR landscape.

**Supplementary Table S3:** Experiments on the antimicrobial resistance prediction tasks.

| Dataset | Baseline Model | LR | | |  | SVM | | |
| --- | --- | --- | --- | --- | --- | --- | --- | --- |
|  |  | F1-score | Recall | Precision |  | F1-score | Recall | Precision |
| HMD-ARG-Class | Ankh | 0.778 | 0.768 | 0.804 |  | 0.891 | 0.877 | 0.932 |
|  | ProteinBERT | 0.875 | 0.847 | 0.941 |  | 0.781 | 0.764 | 0.820 |
|  | ProTrek | 0.844 | 0.837 | 0.862 |  | **0.909** | 0.900 | 0.943 |
|  | ESM-2 | 0.889 | 0.882 | 0.922 |  | 0.844 | 0.839 | 0.860 |
|  | BacLLM (4B) | 0.907 | 0.898 | 0.922 |  | 0.896 | 0.880 | 0.941 |
|  | BacLLM (8B) | **0.919** | 0.908 | 0.941 |  | 0.900 | 0.905 | 0.928 |
| HMD-ARG-Mechanism | Ankh | 0.758 | 0.728 | 0.820 |  | 0.986 | 0.984 | 0.988 |
|  | ProteinBERT | 0.925 | 0.887 | 0.992 |  | 0.915 | 0.872 | 0.990 |
|  | ProTrek | 0.808 | 0.795 | 0.823 |  | 0.960 | 0.937 | 0.989 |
|  | ESM-2 | 0.949 | 0.921 | 0.985 |  | 0.988 | 0.983 | 0.993 |
|  | BacLLM (4B) | **0.984** | 0.981 | 0.988 |  | 0.986 | 0.983 | 0.989 |
|  | BacLLM (8B) | 0.966 | 0.949 | 0.990 |  | **0.988** | 0.984 | 0.993 |
| ARGNet-DB-Class | Ankh | 0.710 | 0.697 | 0.751 |  | 0.917 | 0.917 | 0.930 |
|  | ProteinBERT | 0.885 | 0.872 | 0.909 |  | 0.811 | 0.790 | 0.878 |
|  | ProTrek | 0.826 | 0.814 | 0.848 |  | **0.946** | 0.956 | 0.941 |
|  | ESM-2 | 0.899 | 0.894 | 0.919 |  | 0.867 | 0.858 | 0.893 |
|  | BacLLM (4B) | **0.952** | 0.956 | 0.950 |  | 0.943 | 0.934 | 0.955 |
|  | BacLLM (8B) | 0.937 | 0.947 | 0.931 |  | 0.940 | 0.934 | 0.952 |

**Supplementary Table S4:** Experiments on the Antimicrobial Resistance Prediction task.

| Dataset | Baseline Model | LR |  | | SVM | |
| --- | --- | --- | --- | --- | --- | --- |
|  |  | F1-score | |  | | F1-score |
| ARG-Transferability | Ankh | 0.946 | |  | | 0.971 |
|  | ProteinBERT | 0.965 | |  | | 0.966 |
|  | ProTrek | 0.943 | |  | | **0.979** |
|  | ESM-2 | 0.955 | |  | | 0.959 |
|  | BacLLM (4B) | **0.976** | |  | | 0.978 |
|  | BacLLM (8B) | 0.968 | |  | | 0.976 |
| ARG-noARG | Ankh | 0.869 | |  | | 0.910 |
|  | ProteinBERT | **0.920** | |  | | 0.932 |
|  | ProTrek | 0.880 | |  | | 0.935 |
|  | ESM-2 | 0.897 | |  | | 0.901 |
|  | BacLLM (4B) | 0.914 | |  | | 0.936 |
|  | BacLLM (8B) | 0.916 | |  | | **0.940** |
| BLDB | Ankh | 0.832 | |  | | 0.830 |
|  | ProteinBERT | 0.978 | |  | | 0.974 |
|  | ProTrek | 0.980 | |  | | **1.000** |
|  | ESM-2 | 0.980 | |  | | 0.828 |
|  | BacLLM (4B) | **1.000** | |  | | **1.000** |
|  | BacLLM (8B) | **1.000** | |  | | **1.000** |

**Supplementary Table S5:** Cosine similarities between protein representations and TolC in the TolC-centered protein association network.

| Gene ID | Similarity |
| --- | --- |
| KPN_00995 | 0.7732 |
| KPN_03299 | 0.6608 |
| acrB | 0.6548 |
| acrF | 0.6388 |
| KPN_02970 | 0.6230 |
| KPN_01486 | 0.6189 |
| KPN_00561 | 0.6161 |
| acrD | 0.6075 |
| yohG | 0.6028 |
| KPN_02064 | 0.6021 |
| tolQ | 0.5989 |
| KPN_00560 | 0.5985 |
| KPN_04041 | 0.5770 |
| tolA | 0.5693 |
| ydhK | 0.5668 |
| KPN_02144 | 0.5651 |
| KPN_04040 | 0.5628 |
| acrE | 0.5590 |
| acrA | 0.5522 |

**Note:** For convenience, proteins are labeled using their corresponding gene symbols or locus tags. The network was constructed from BacLLM-derived protein sequence embeddings and represents protein-level associations rather than gene-level interaction**s.**

**Supplementary Table S6:** In the *Klebsiella pneumoniae* dataset, clustering was performed using a 90% sequence similarity threshold, and LR was employed as the classifier in downstream tasks.

| K | ESM2 | Ankh | ProteinBERT | ProTrek | BacLLM(8B) | BacLLM(4B) |
| --- | --- | --- | --- | --- | --- | --- |
| 1 | 0.4434 | 0.4434 | 0.4434 | 0.2827 | 0.4434 | 0.4434 |
| 2 | 0.5342 | 0.5525 | 0.4982 | 0.4037 | 0.5434 | 0.5434 |
| 3 | 0.5647 | 0.5827 | 0.5644 | 0.5799 | 0.5799 | 0.5647 |
| 4 | 0.6240 | 0.6071 | 0.6101 | 0.6240 | 0.6165 | 0.6287 |
| 5 | 0.6402 | 0.6254 | 0.6524 | 0.6402 | 0.6565 | 0.6565 |
| 6 | 0.6524 | 0.6406 | 0.6524 | 0.6402 | 0.6849 | 0.6849 |
| 7 | 0.6646 | 0.6707 | 0.6524 | 0.6402 | 0.6849 | 0.6849 |
| 8 | 0.6768 | 0.6768 | 0.6890 | 0.6463 | 0.6971 | 0.6971 |
| 9 | 0.6951 | 0.6768 | 0.6890 | 0.6646 | 0.6971 | 0.6971 |
| 10 | 0.7012 | 0.6890 | 0.7256 | 0.6707 | 0.6971 | 0.6971 |

**Supplementary Table S7:** In the *Klebsiella pneumoniae* dataset, clustering was performed using a 90% sequence similarity threshold, and SVM was employed as the classifier in downstream tasks.

| K | ESM2 | Ankh | ProteinBERT | ProTrek | BacLLM(8B) | BacLLM(4B) |
| --- | --- | --- | --- | --- | --- | --- |
| 1 | 0.3162 | 0.3815 | 0.3071 | 0.4066 | 0.4386 | 0.3904 |
| 2 | 0.3796 | 0.4943 | 0.3772 | 0.4639 | 0.5067 | 0.4615 |
| 3 | 0.4172 | 0.5552 | 0.4101 | 0.5188 | 0.5676 | 0.5189 |
| 4 | 0.4599 | 0.5983 | 0.4345 | 0.5358 | 0.6131 | 0.5328 |
| 5 | 0.4660 | 0.6288 | 0.4528 | 0.5480 | 0.6314 | 0.5819 |
| 6 | 0.4782 | 0.6337 | 0.4772 | 0.5602 | 0.6636 | 0.6567 |
| 7 | 0.5042 | 0.6337 | 0.4873 | 0.5620 | 0.6916 | 0.6628 |
| 8 | 0.5073 | 0.6337 | 0.4873 | 0.5888 | 0.7044 | 0.6668 |
| 9 | 0.5073 | 0.6337 | 0.4904 | 0.6010 | 0.7044 | 0.6906 |
| 10 | 0.5090 | 0.6337 | 0.4985 | 0.6010 | 0.7170 | 0.7028 |

**Supplementary Table S8:** In the *Klebsiella pneumoniae* dataset, clustering was performed using a 95% sequence similarity threshold, and LR was employed as the classifier in downstream tasks.

| K | ESM2 | Ankh | ProteinBERT | ProTrek | BacLLM(8B) | BacLLM(4B) |
| --- | --- | --- | --- | --- | --- | --- |
| 1 | 0.4269 | 0.4269 | 0.4269 | 0.2724 | 0.4269 | 0.4269 |
| 2 | 0.5177 | 0.5346 | 0.4831 | 0.3913 | 0.5262 | 0.5262 |
| 3 | 0.5458 | 0.5720 | 0.5552 | 0.5608 | 0.5608 | 0.5458 |
| 4 | 0.6110 | 0.5945 | 0.5983 | 0.6110 | 0.6058 | 0.6170 |
| 5 | 0.6260 | 0.6114 | 0.6372 | 0.6260 | 0.6410 | 0.6410 |
| 6 | 0.6372 | 0.6264 | 0.6372 | 0.6260 | 0.6672 | 0.6672 |
| 7 | 0.6485 | 0.6541 | 0.6372 | 0.6260 | 0.6672 | 0.6672 |
| 8 | 0.6597 | 0.6597 | 0.6709 | 0.6316 | 0.6784 | 0.6784 |
| 9 | 0.6822 | 0.6597 | 0.6709 | 0.6485 | 0.6784 | 0.6784 |
| 10 | 0.6878 | 0.6709 | 0.7159 | 0.6541 | 0.6784 | 0.6784 |

**Supplementary Table S9:** In the *Klebsiella pneumoniae* dataset, clustering was performed using a 95% sequence similarity threshold, and SVM was employed as the classifier in downstream tasks.

| K | ESM2 | Ankh | ProteinBERT | ProTrek | BacLLM(8B) | BacLLM(4B) |
| --- | --- | --- | --- | --- | --- | --- |
| 1 | 0.2986 | 0.4082 | 0.3014 | 0.4185 | 0.4569 | 0.4335 |
| 2 | 0.3436 | 0.4897 | 0.3436 | 0.4831 | 0.5252 | 0.5102 |
| 3 | 0.3754 | 0.5514 | 0.3661 | 0.5449 | 0.5926 | 0.5608 |
| 4 | 0.4260 | 0.5945 | 0.3885 | 0.5617 | 0.6441 | 0.5777 |
| 5 | 0.4372 | 0.6245 | 0.4054 | 0.5767 | 0.6610 | 0.6132 |
| 6 | 0.4597 | 0.6301 | 0.4279 | 0.6048 | 0.6928 | 0.6822 |
| 7 | 0.4747 | 0.6376 | 0.4316 | 0.6048 | 0.7172 | 0.6990 |
| 8 | 0.5149 | 0.6376 | 0.4466 | 0.6273 | 0.7284 | 0.7028 |
| 9 | 0.5187 | 0.6376 | 0.4625 | 0.6385 | 0.7322 | 0.7262 |
| 10 | 0.5187 | 0.6488 | 0.4737 | 0.6385 | 0.7359 | 0.7374 |

**Supplementary Table S10:** In the *Klebsiella pneumoniae* dataset, clustering was performed using a 99.5% sequence similarity threshold, and LR was employed as the classifier in downstream tasks.

| K | ESM2 | Ankh | ProteinBERT | ProTrek | BacLLM(8B) | BacLLM(4B) |
| --- | --- | --- | --- | --- | --- | --- |
| 1 | 0.4089 | 0.4089 | 0.4089 | 0.2716 | 0.4089 | 0.4089 |
| 2 | 0.4922 | 0.5108 | 0.4552 | 0.3858 | 0.5108 | 0.5108 |
| 3 | 0.5200 | 0.5478 | 0.5293 | 0.5339 | 0.5339 | 0.5200 |
| 4 | 0.5848 | 0.5663 | 0.5756 | 0.5848 | 0.6033 | 0.6126 |
| 5 | 0.6033 | 0.5848 | 0.6126 | 0.6126 | 0.6404 | 0.6466 |
| 6 | 0.6126 | 0.6033 | 0.6126 | 0.6126 | 0.6743 | 0.6743 |
| 7 | 0.6219 | 0.6311 | 0.6126 | 0.6126 | 0.6836 | 0.6836 |
| 8 | 0.6404 | 0.6404 | 0.6496 | 0.6172 | 0.6929 | 0.6929 |
| 9 | 0.6682 | 0.6404 | 0.6496 | 0.6404 | 0.6929 | 0.6929 |
| 10 | 0.6682 | 0.6496 | 0.6867 | 0.6404 | 0.6929 | 0.6929 |

**Supplementary Table S11:** In the *Klebsiella pneumoniae* dataset, clustering was performed using a 99.5% sequence similarity threshold, and SVM was employed as the classifier in downstream tasks.

| K | ESM2 | Ankh | ProteinBERT | ProTrek | BacLLM(8B) | BacLLM(4B) |
| --- | --- | --- | --- | --- | --- | --- |
| 1 | 0.2870 | 0.4521 | 0.2777 | 0.5324 | 0.5509 | 0.5293 |
| 2 | 0.3425 | 0.5354 | 0.3055 | 0.5694 | 0.6157 | 0.5895 |
| 3 | 0.4027 | 0.5910 | 0.3287 | 0.5972 | 0.6620 | 0.6404 |
| 4 | 0.4537 | 0.6342 | 0.3657 | 0.6250 | 0.6990 | 0.6496 |
| 5 | 0.4722 | 0.6712 | 0.3919 | 0.6466 | 0.7175 | 0.6728 |
| 6 | 0.4953 | 0.6712 | 0.4243 | 0.6651 | 0.7407 | 0.7237 |
| 7 | 0.5046 | 0.6774 | 0.4336 | 0.6743 | 0.7530 | 0.7469 |
| 8 | 0.5416 | 0.6774 | 0.4429 | 0.6929 | 0.7685 | 0.7469 |
| 9 | 0.5509 | 0.6774 | 0.4552 | 0.7021 | 0.7685 | 0.7592 |
| 10 | 0.5694 | 0.6867 | 0.4645 | 0.7021 | 0.7731 | 0.7685 |

**Supplementary Table S12:** In the *Escherichia* dataset, experiments were repeated using 20 random seeds, and the results were averaged across runs. LR was employed as the classifier in downstream tasks.

| K | ESM2 | Ankh | ProteinBERT | ProTrek | BacLLM(8B) | BacLLM(4B) |
| --- | --- | --- | --- | --- | --- | --- |
| 1 | 0.6072 | 0.6072 | 0.6072 | 0.6072 | 0.6666 | 0.6666 |
| 2 | 0.6743 | 0.6635 | 0.6635 | 0.6743 | 0.6743 | 0.6743 |
| 3 | 0.7988 | 0.8002 | 0.7382 | 0.7989 | 0.7949 | 0.7930 |
| 4 | 0.8480 | 0.8864 | 0.8340 | 0.8397 | 0.8665 | 0.8665 |
| 5 | 0.8863 | 0.8934 | 0.8889 | 0.8934 | 0.9175 | 0.9175 |
| 6 | 0.9061 | 0.9132 | 0.9094 | 0.9068 | 0.9347 | 0.9354 |
| 7 | 0.9246 | 0.9285 | 0.9157 | 0.9183 | 0.9482 | 0.9482 |
| 8 | 0.9348 | 0.9355 | 0.9278 | 0.9304 | 0.9577 | 0.9577 |
| 9 | 0.9380 | 0.9418 | 0.9361 | 0.9431 | 0.9590 | 0.9590 |
| 10 | 0.9437 | 0.9489 | 0.9495 | 0.9482 | 0.9597 | 0.9590 |

**Supplementary Table S13:** In the *Escherichia* dataset, experiments were repeated using 20 random seeds, and the results were averaged across runs. SVM was employed as the classifier in downstream tasks.

| K | ESM2 | Ankh | ProteinBERT | ProTrek | BacLLM(8B) | BacLLM(4B) |
| --- | --- | --- | --- | --- | --- | --- |
| 1 | 0.6072 | 0.6072 | 0.6072 | 0.6679 | 0.6653 | 0.6603 |
| 2 | 0.6647 | 0.6743 | 0.6635 | 0.7019 | 0.7695 | 0.6986 |
| 3 | 0.6852 | 0.6852 | 0.6852 | 0.8028 | 0.8176 | 0.8033 |
| 4 | 0.8089 | 0.8121 | 0.8020 | 0.8447 | 0.8640 | 0.8621 |
| 5 | 0.8089 | 0.8153 | 0.8020 | 0.8645 | 0.8979 | 0.9016 |
| 6 | 0.8153 | 0.8153 | 0.8153 | 0.9100 | 0.9080 | 0.9125 |
| 7 | 0.8952 | 0.8952 | 0.8952 | 0.9214 | 0.9189 | 0.9310 |
| 8 | 0.9220 | 0.9220 | 0.9373 | 0.9342 | 0.9336 | 0.9380 |
| 9 | 0.9437 | 0.9412 | 0.9482 | 0.9418 | 0.9381 | 0.9432 |
| 10 | 0.9520 | 0.9488 | 0.9577 | 0.9450 | 0.9425 | 0.9451 |

**Supplementary Table S14: Parameter ablation experiments of the BacLLM(4B) model.**

|  | G(θ) | Text Augmentation | F1-Score |
| --- | --- | --- | --- |
| SVM | **√** | √ | **0.943** |
|  | √ | × | 0.894 |
|  | × | √ | 0.935 |
|  | × | × | 0.903 |
| LR | √ | √ | **0.952** |
|  | √ | × | 0.933 |
|  | × | √ | 0.948 |
|  | × | × | 0.950 |

Note: G(θ) denotes the sequence semantic enhancement module, while Text Augmentation refers to the text semantic enhancement module. This experiment conducts an ablation study on BacLLM (4B).

**Supplementary Table S15. Parameter ablation experiments of the BacLLM(8B) model.**

|  | G(θ) | Text Augmentation | F1-Score |
| --- | --- | --- | --- |
| SVM | **√** | √ | **0.940** |
|  | √ | × | 0.897 |
|  | × | √ | 0.901 |
|  | × | × | 0.900 |
| LR | √ | √ | **0.937** |
|  | √ | × | 0.934 |
|  | × | √ | 0.934 |
|  | × | × | 0.936 |

Note: G(θ) denotes the sequence semantic enhancement module, while Text Augmentation refers to the text semantic enhancement module. This experiment conducts an ablation study on BacLLM (8B).
